# High-throughput cultivation and isolation of environmental anaerobes using selectively permeable hydrogel capsules

**DOI:** 10.1101/2025.05.06.652511

**Authors:** Hugo Sallet, Marion Calvo, Matteo Titus, Nicolas Jacquemin, Karin Lederballe Meibom, Rizlan Bernier-Latmani

## Abstract

Over the past two decades, metagenomics has greatly expanded our understanding of microbial phylogenetic and metabolic diversity. However, most microbial taxa remain uncultured, hindering research and biotechnological applications. Isolating environmental anaerobes using traditional methods is particularly cumbersome and low throughput. Here, we present a novel, high-throughput approach for the cultivation and isolation of anaerobes, which involves trapping and growing single microbes within selectively permeable hydrogel capsules followed by fluorescence-activated cell sorting to distribute compartmentalized isolates into liquid medium for further growth. We show that diverse anaerobes can grow within capsules and that slower-growing ones (e.g. methanogens) can be enriched with this platform. We also applied this approach to isolate anaerobes from soil, including strains of the sulfate-reducing bacteria *Desulfovibrio desulfuricans* and *Nitratidesulfovibrio vulgaris*. Overall, this work introduces a robust, high-throughput alternative to traditional techniques for isolating environmental anaerobes and expands the emerging set of microfluidics-based tools for the cultivation of novel taxa.

## Introduction

While cultivation-independent methods are now a gold standard for exploring microbiomes, the isolation of microorganisms remains imperative for studying their physiology, ecology, and evolution [1]. Cultivating individual strains in the laboratory offers numerous advantages: specific metabolisms (e.g. pollutant degradation, antibiotic synthesis) can be exploited, hypotheses (e.g. genome-based predictions) can be tested experimentally under controlled conditions, and high-quality genomes can be obtained to expand and enhance current databases.

However, our ability to isolate and cultivate microorganisms is extremely limited: it is estimated that less than 3% of prokaryotic species have been successfully cultivated [2–4], and around 90% of these are affiliated with only four phyla (Pseudomonadota, Bacillota, Actinomycetota, and Bacteroidota) [5,6]. These numbers vary considerably across habitats: while cultured taxa predominate in human-associated microbiomes, most taxa found in underexplored environments (e.g. soil, terrestrial subsurface, hydrothermal vents) remain uncultured [7]. The discrepancy between the number of microorganisms observed under a microscope and those cultured using traditional methods (typically, agar plating), known as the ‘Great Plate Count Anomaly’ [8], highlights our inability to mimic the complex conditions necessary for microbial growth.

Growth requirements can vary largely across taxa, and include not only suitable environmental conditions (e.g. temperature, pH, redox conditions) but also specific growth factors which are generally unknown [1]. In fact, even when these conditions are fulfilled, some microorganisms may not be able to grow because they are inhibited by others (e.g. competition for nutrients) or, in contrast, because they need others to grow (e.g. syntrophy, episymbiosis) [1]. Finally, conventional culture media often contain high concentrations of organic substrates and thus favour fast-growing taxa, which challenges the efforts to enrich and isolate slow-growers (e.g. oligotrophs) [9].

Efforts to improve ‘cultivability’ rates by modifying artificial nutrient media have met limited success [10–12]. Culturomics approaches, which combine the diversification of culture conditions with systematic isolate screening (e.g. MALDI-TOF, colony imaging), have allowed the isolation of hundreds of novel taxa [13–15], but remain laborious and have only scarcely been applied to non-human microbiomes. Additionally, such approaches based on agar plating are ill-suited to cultivate environmental anaerobes [16,17], some of which are key isolation targets (e.g. methanogens, anammox bacteria) [1,18]. In particular, some obligate anaerobes (i.e. unable to utilize oxygen as a terminal electron acceptor) grow poorly at gas-liquid interfaces and require specific gases as energy or carbon sources. Alternative approaches, such as the roll-tube and the dilution-to-extinction methods [17,19–21], have been employed to cultivate anaerobes but they are time-consuming and do not lend themselves to screening large numbers of isolates.

Within the past two decades, microfluidics-based methods have been developed to raise the throughput rate and facilitate the cultivation of low-abundance taxa. For instance, Ma *et al.* employed a custom microfluidic device (SlipChip) to concurrently perform genetic screening and cultivation of isolates in a high-throughput manner [22], enabling the targeted isolation of a novel anaerobic bacterium (member of Ruminococcaceae) from the human gut [23]. Another notable example is the iChip platform designed as an array of miniature diffusion chambers (microcompartments filled with microbial inoculum in agar-based medium and sealed with porous membranes), which advantageously allows for *in situ* cultivation, thus providing organisms with key environmental growth factors [24]. Although these platforms have aided the discovery of novel taxa, they have been used rarely since, possibly because of the difficulty to reliably craft and manipulate such miniature devices.

On the other hand, the increasing availability of fluorescence-activated cell sorting (FACS) has encouraged the development of isolation strategies based on trapping and growing single microorganisms in picolitre-size compartments, sorting these compartments based on the biomass enclosed within, and distributing these individually in medium to further grow the isolates. For instance, Zengler *et al*. have isolated novel strains from seawater and soil by cultivating microorganisms within agarose beads (i.e. permeable gel compartments), sorting and re-growing in microwell plates [25]. Recently, McCully *et al.* used a double-emulsion (i.e. water-in-oil-in-water) platform to grow stool-derived microorganisms. They observed an enrichment of slow-growing taxa (e.g. the Negativicutes *Phascolarctobacterium faecium*) – which they attributed to limited competition across compartments stemming from the low water-oil permeability [26] – though none of these taxa could be isolated using FACS.

Over the past decade, hydrogel capsules, i.e., microcompartments comprising a liquid core and a solid shell, have been increasingly popular in drug delivery and single-cell assays due to their robustness, biocompatibility and selective permeability [27–29]. Additionally, microfluidic instruments and reagent kits for capsule generation have become commercially available [30], making this technology accessible to microbiologists with no prior technical experience. Yet, although capsules have been shown to be compatible with microbial growth, they have not been used for isolating microorganisms.

This proof-of-concept study demonstrates the potential of selectively permeable hydrogel capsules (hereafter capsules) as a platform for the cultivation and isolation of anaerobes. We show that capsules support the growth of diverse anaerobic taxa, including those which fail to form colonies on agar. We also found that soil microorganisms grown in capsules were more diverse than those grown on agar and within agarose beads, suggesting that capsules may be advantageous to isolate novel taxa. Finally, following in-capsule cultivation, FACS was used to isolate anaerobes (including sulfate-reducing bacteria, SRB) from soil in a high-throughput manner. This approach can be adapted to different sample types and growth conditions and may be compatible with *in situ* cultivation.

## Materials and methods

In this work, we trap single microorganisms in capsules and grow them within. First, we grow known isolates anaerobically within capsules to establish feasibility. Next, we do the same with soil-extracted microorganisms, which were also grown simultaneously in other cultivation platforms (liquid, agar, agarose beads, water-in-oil droplets) to compare the cultivation potential of the different methods. We also follow the temporal changes in a soil-derived, capsule-entrapped microbial community. Finally, following in-capsule cultivation, we use FACS to isolate anaerobes – either directly extracted from soil or from enrichment cultures. See summary in Fig. S1.

### Generation of capsules

Capsules were produced with the Onyx instrument using the SPC Innovator Kit (Atrandi Biosciences, Vilnius, Lithuania) [31]. Capsules are aqueous two-phase systems (ATPS) consisting of two immiscible aqueous phases, a core polymer and a shell polymer, combined in a microfluidic chip (Fig. 1A). While the former remains in the capsule core, the latter instantaneously migrates outwards upon encapsulation. Microbial cells were added to the core polymer immediately prior to encapsulation. Flow rates of 35 µl/h and 250 µl/h were used for the aqueous and oil phase, respectively. The encapsulation was monitored in real time with the high-speed camera of the Onyx instrument (Movie 1). The emulsion was briefly exposed to 405-nm light to induce crosslinking of the shell polymer, and subsequently broken as indicated in the supplier’s protocol [31]. This results in an aqueous suspension of 30-40 µm (14-33 pl) capsules (coefficient of variation ∼ 15%), which are permeable to molecules smaller than 160 kDa (Fig. 1B). To maximize the encapsulation of single cells, the cell density in the aqueous phase was adjusted to 10^7^ cells/ml, achieving a Poisson parameter (λ) of 0.3. All steps were performed under an anoxic atmosphere (Fig. S2).

**Figure 1:**
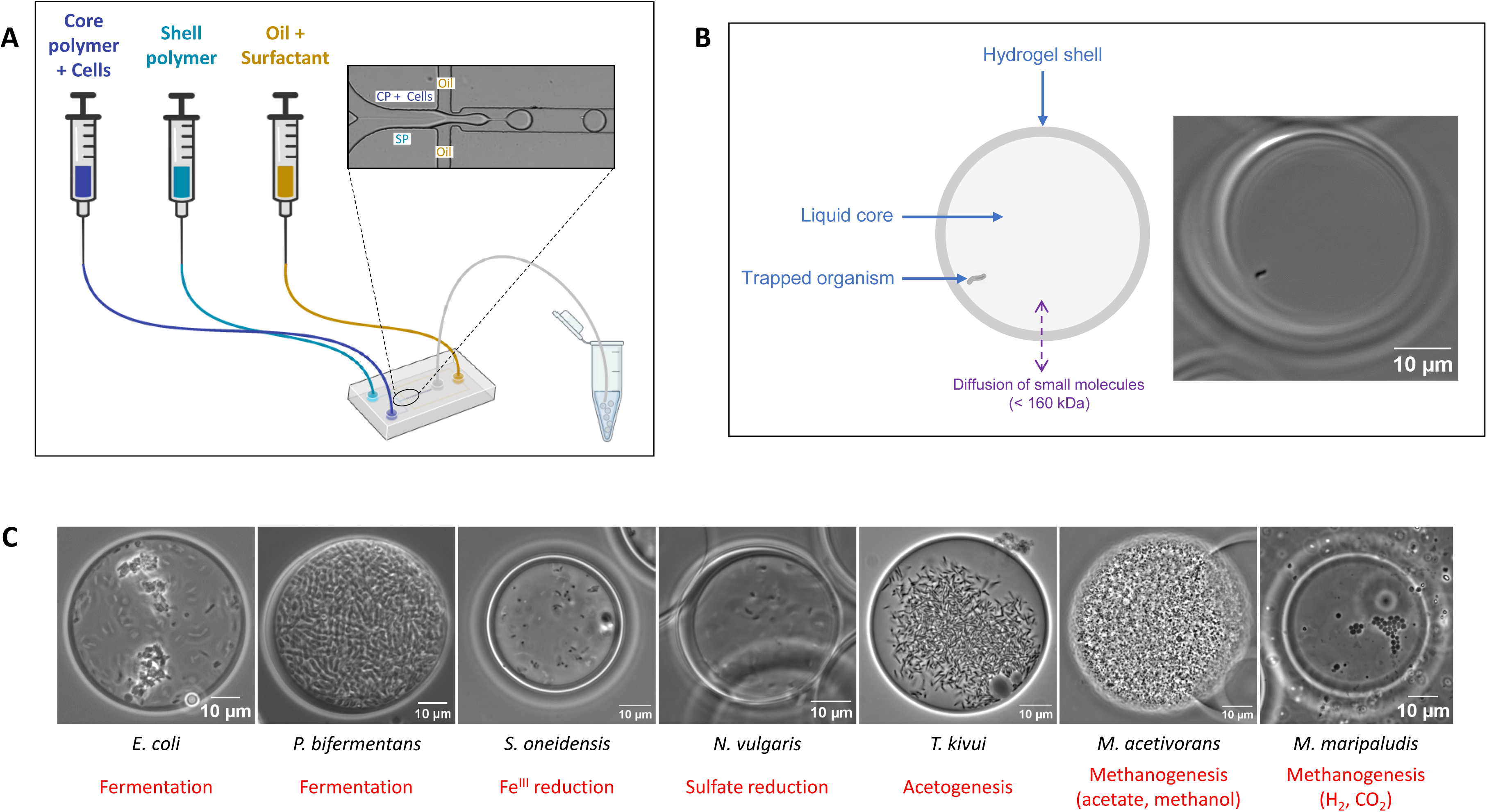
Single-cell encapsulation and growth of anaerobic microorganisms within capsules. (A) Overview of the capsule generation process (inset: microscope image of microfluidic chip during capsule generation). (B) Schematic and microscopy image showing a capsule enclosing a single bacterium (*Nitratidesulfovibrio vulgaris*). (C) Microscopy images of anaerobic strains growing within capsules. From left to right: *Escherichia coli* TB205 (18 h incubation), *Paraclostridium bifermentans* EML (24 h incubation), *Shewanella oneidensis* MR-1 (24 h incubation), *Nitratidesulfovibrio vulgaris* (2-day incubation), *Thermoanaerobacter kivui* (24 h incubation), *Methanosarcina acetivorans* C2A (18-day incubation) and *Methanococcus maripaludis* MM901 (5-day incubation). The type of anaerobic metabolism used by the strains is specified in red below each image. CP: core polymer, SR: shell polymer.

### Cultivation of known anaerobic strains in capsules

Seven phylogenetically and metabolically diverse anaerobic strains (Table 1) were individually compartmentalized into capsules, starting from fresh cultures. Following emulsion breaking, the capsules were resuspended in the appropriate media (Table 1) and incubated under anoxic conditions. Capsules were imaged under a microscope (Eclipse Ni-E, Nikon, Tokyo, Japan). Fluorescence imaging was also performed following a brief staining protocol (10-min incubation, 1 µM SYTO-9).

**Table 1:** Characteristics and cultivation conditions of microbial strains used in this study.

| Strain name | Source | Obligate/Facultative anaerobe | Metabolic group | Cultivation conditions<sup>a</sup> |
| --- | --- | --- | --- | --- |
| *Paraclostridium bifermentans* EML | Isolated from paddy soil [51] | Facultative | Fermenter | LB medium (BD Difco™), 30°C |
| *Shewanella oneidensis* MR-1 | Lab collection | Facultative | Iron-reducing bacterium | M9 medium [52] amended with sodium lactate (20 mM) and ferric citrate (40 mM), 30°C |
| *Escherichia coli* TB205 (engineered by Bergmiller et al. [53]) | Gift from Tobias Bergmiller | Facultative | Fermenter | LB medium (BD Difco™), 37°C |
| *Methanosarcina acetivorans* C2A | DSMZ (strain 2834) | Obligate | Methanogen | DSMZ 141c medium, N<sub>2</sub>/CO<sub>2</sub> (80/20 v/v) headspace, 37°C |
| *Nitratidesulfovibrio vulgaris* | Lab collection | Obligate | Sulfate-reducing bacterium | DSMZ medium 63, 37°C |
| *Thermoanaerobacter kivui* | DSMZ (strain 2030) | Obligate | Homoacetogen | Complex medium described by Basen et al. [54], H<sub>2</sub>/CO<sub>2</sub> (80/20 v/v) headspace, 60°C |
| *Methanococcus maripaludis* MM901 (engineered by Costa et al. [55]) | Gift from Wenyu Gu | Obligate | Methanogen | DSMZ 141c medium, H<sub>2</sub>/CO<sub>2</sub> (80/20 v/v) headspace, 37°C |

<sup>a</sup> Media were brought to a boil, cooled down to room temperature under a gas flow (100% N<sub>2</sub> or 80%:20% N<sub>2</sub>:CO<sub>2</sub>), and dispensed into 200-ml serum bottles (100 ml of medium per bottle) under the same gas atmosphere. The bottles were sealed with butyl rubber stoppers and aluminium crimps and autoclaved at 121°C for 30 min. Prior to autoclaving, the headspace of the serum bottles was flushed with the recommended gas.

### Soil sampling and extraction of soil microorganisms

Soil (upper 10 cm) was sampled from two adjacent rice paddies (A and B) in Mont-Vully, Switzerland (46°58’06.5”N 7°03’39.2”E), which were continuously flooded throughout the growing season (4 months). Samples were collected either 2 months after the harvest (A_AH_ and B_AH_) or shortly before the following year’s harvest (A_BH_ and B_BH_). Immediately after collection, the samples were placed in Mylar bags filled with argon and stored at 4°C. An aliquot of soil A_AH_ was stored at -80°C for DNA extraction. Soil physicochemical characteristics are described in Table S1.

To extract microorganisms, soil was mixed (1:4 volume) with a sodium pyrophosphate solution (0.2% in PBS) by vortexing and blending (30 s^-1^, 2 min) with a MM 400 mixer mill (Retsch, Haan, Germany). The resulting slurry was left to decant (30 min) and the supernatant was transferred onto a Nycodenz (Serumwerk Bernburg, Bernburg, Germany) solution (80% in PBS) at 4:1 volume ratio. After centrifugation (15 000 g for 90 min at 4°C), the cell-containing fraction was collected, washed several times in PBS and filtered (10 µm).

### Comparison of cultivation platforms for soil-derived microorganisms

Soil-extracted microorganisms were grown in capsules, in agarose beads, in water-in-oil droplets, in liquid, and on agar. Water-in-oil droplets (hereafter droplets) were generated with the Onyx platform using the Droplet Generation Kit (Atrandi Biosciences), with an aqueous phase composed of microbial cells suspended in medium. Agarose beads were generated as described by Schaerli [32]. For droplets and agarose beads, flow rates of 300 µl/h and 700 µl/h were used for the aqueous and oil phases, respectively. Cell density was adjusted as described above. Soil microorganisms were also spread on agar plates or directly inoculated in liquid medium. All steps were performed under an anoxic atmosphere and all cultivation platforms used the same Minimal Soil Medium (MSM10, see SI).

After incubation, microbial cells were extracted from capsules by dissolving the shell polymer with the Release Reagent (Atrandi Biosciences) at RT for 1 h under agitation (500 rpm). Agarose beads were dissolved by incubating with β-agarase I (5 units, 42°C, 30 min), following a preheating step (65°C for 10 min). Capsules and agarose beads were washed twice in PBS before the extraction. Microorganisms were released from droplets by breaking the emulsion with 1*H*,1*H*,2*H*,2*H*-Perfluoro-1-octanol, as described previously [33]. Microbes grown on agar were harvested by adding PBS (2 ml) to agar plates, scraping the surface with an L-shaped spreader, and collecting the wash solution. Cells were collected from the liquid culture by centrifugation (8 000 g, 5 min).

### Temporal change in composition of capsule-entrapped microbial community

Soil microorganisms were individually trapped in capsules and incubated in MSM10 medium for varying durations (24, 48, 72 and 144 hours) to evaluate the potential enrichment in slower-growing strains over time.

### Isolation of slow-growing soil anaerobes

The isolation of anaerobes was first attempted by encapsulating soil-extracted microorganisms and incubating the capsules either in MSM or in 10-ml serum bottles filled with enrichment media (DSMZ 135, DSMZ 311, or DSMZ 141c). Isolation was also attempted by encapsulating microorganisms from soil-derived cultures enriched in SRB and methanogens, using modified Postgate B and modified DSMZ 141c, respectively, as enrichment media. See SI for details. After growth, capsules were separated using FACS. Capsules were sorted with a FACSAria Fusion (BD Biosciences, Franklin States, NJ, USA), see details in SI. Prior to sorting, the capsules were washed three times in PBS and stained with SYTO-9 (2 µM, 15-min incubation), a fluorescent dye retaining cell viability. Two controls were prepared to facilitate the gating: one without SYTO-9 and another with SYTO-9 but incubated in PBS. More details on the gating are shown in Fig. S3. Single capsules were distributed into individual wells of 96-well plates filled with medium (200 µl/well) and incubated at 30°C in an anoxic chamber, or in a nox.18 anaerobic jar (SY-LAB, Purkersdorf, Austria) filled with H_2_ (80%) and CO_2_ (20%).

### DNA extraction and sequencing

The DNA from the soil-derived cultures was extracted with the PowerSoil Pro kit (Qiagen, Hilden, Germany) and concentrations were measured with a Qubit fluorometer (Invitrogen, Waltham, MA, USA). These samples were standardized to the same DNA concentration. Library preparation (Nextera DNA Flex, Illumina, San Diego, CA, USA) and sequencing (Novaseq 6000, Illumina, or Aviti, Element Biosciences) were performed at the Lausanne Genomic Technologies Facility (University of Lausanne, Switzerland). The taxonomy of soil A_AH_ was profiled by full-length 16S rRNA gene sequencing, performed on a Sequel II (Pacific Biosciences, Menlo Park, CA, USA).

### Bioinformatic analysis

Shotgun metagenomic data were analyzed to profile the taxonomy of the cultures. Paired-end reads underwent quality control (fastp) to remove adapters and low-quality reads, followed by an additional quality check with FASTQC. High-quality reads from individual samples were then independently assembled into contigs with MEGAHIT. Binning of assembled contigs was performed with BASALT, grouping contigs from all samples into one set of dereplicated metagenome-assembled genomes (MAGs). MAG quality was evaluated with Quast, and CheckM2 was used to further assess completeness and contamination. Only MAGs with >50% completeness and <10% contamination were retained for downstream analysis. MAGs were taxonomically classified with GTDB-Tk default database. See SI for details.

### Taxonomic identification of isolates

Cell lysis was performed by mixing (1:4 v/v) the pure cultures with a Triton-X-100 buffer (0.1% v/v in TE) and heating at 99°C for 5 min. The lysates were spun down and the supernatant was used directly as DNA template in PCR reactions to amplify the near full-length 16S rRNA genes, using 27F (5’-AGA GTT TGA TCC TGG CTC AG-3’) and 1492R (5’-GGT TAC CTT GTT ACG ACT T-3’) primers. Further sample purification and Sanger sequencing were performed by Microsynth (Balgach, Switzerland).

## Results

Selectively permeable capsules are hydrogel microcompartments whose pore size is both small enough to retain cells and large enough to allow the diffusion of small biomolecules (Fig. 1B). In addition, they include a rigid shell, allowing physical separation from bulk solution, and a liquid core, allowing planktonic microbial growth, making them distinct from other types of microcompartments (droplets, agarose beads) (Fig. S4). We set out to leverage the recent commercial availability of this technology to provide a proof-of-concept for its use to cultivate and isolate environmental anaerobes.

### Capsules support the growth of known anaerobic strains

Seven anaerobic strains, varying in phylogeny, metabolism, and oxygen sensitivity (Table 1), were all able to grow within capsules (Fig. 1C). Fluorescence microscopy could successfully reveal microbial cells enclosed within capsules (Fig. S5), and their active metabolic state was evidenced by their motility (Movies 2-6). Interestingly, both a planktonic lifestyle and biofilm formation were observed within capsules containing *E. coli* (Fig. 1C, Movie 3). Therefore, diverse microbial species can grow anaerobically within capsules.

### Growth of distinct soil microbial taxa across cultivation platforms

We then examined whether soil microorganisms could grow anaerobically within capsules, and, if so, which taxa would preferentially grow in this cultivation platform compared to others, i.e. liquid, agar, agarose beads, droplets. The latter two represent compartmentalized platforms with characteristics distinct from capsules (Fig. S4).

The soil was dominated by microbial taxa in the phyla Bacillota (formerly Firmicutes), Pseudomonadota (formerly Proteobacteria) and Bacteroidota, and also contained members of Desulfobacterota (expected SRB) in substantial (6%) proportions (Fig. S6).

Soil-extracted microbial cells were cultured individually in compartments (capsules, agarose beads, droplets), and concomitantly on solid (agar) and in liquid medium (Fig. 2A) and growth was observed microscopically within a few days (Fig. S7). After one week of growth, the highest richness (number of MAGs) was observed in the liquid culture (155 ± 2.6 MAGs), and the lowest on agar (57 ± 3.6 MAGs) (Fig. 2B). Within compartmentalized systems, capsules (131 ± 4.4 MAGs) showed higher richness than droplets (91 ± 4.0 MAGs) and agarose beads (85 ± 14.6 MAGs) (Fig. 2B). A strong deviation from the native community was observed in the agar, agarose beads, and liquid cultures (Fig. 2C), whose resulting communities were associated with 8, 7 and 15 unique signature MAGs, respectively (Fig. 2D). As expected, agar plating did not allow the growth of most microbial taxa (including 58 MAGs found in all other conditions) (Fig. 2D), and strongly favoured phyla associated with faster growth rates and aerobic metabolisms (Actinomycetota, Bacillota and Pseudomonadota) (Fig. 2E). On the other hand, capsule-grown microorganisms were more reflective of the original community structure and showed high similarity with liquid cultures (22 shared MAGs) (Fig. 2C-D). Notably, phyla associated with slow-growing taxa (Methanobacteriota, Halobacteriota, Fusobacteriota) were all found in liquid and capsules, and to a lesser extent in agarose beads and emulsion cultures, but none were detected in the agar cultures (Fig. 2E). In particular, seven taxa in the classes Methanobacteria (bin111, bin54, bin105, bin69, bin104) and Methanosarcinia (bin170, bin57) were identified as methanogens, and were present in the capsules (Fig. 2E, Table S2). Interestingly, poorly characterized taxa, such as *Anaerorhabdus* sp. (bin 162) and Fusobacteriaceae gen. sp. (bin 148), were only detected in liquid and capsules (Fig. 2E, Table S2). We also performed a similar analysis directly on the short-read sequences (using mOTUs) – to exclude potential biases caused by metagenomic assembly – and observed similar trends (Fig. S8). Interestingly, microorganisms within the class Clostridia were only shown to grow in liquid, agarose beads, and capsules (Fig. S9-10). The mOTUs taxonomy also showed evidence of Desulfovibrionia taxa (putative SRB) growing in liquid and within capsules (Fig. S10). Collectively, these results indicate that a broad range of soil microorganisms, including poorly characterized taxa, can grow within capsules.

**Figure 2:**
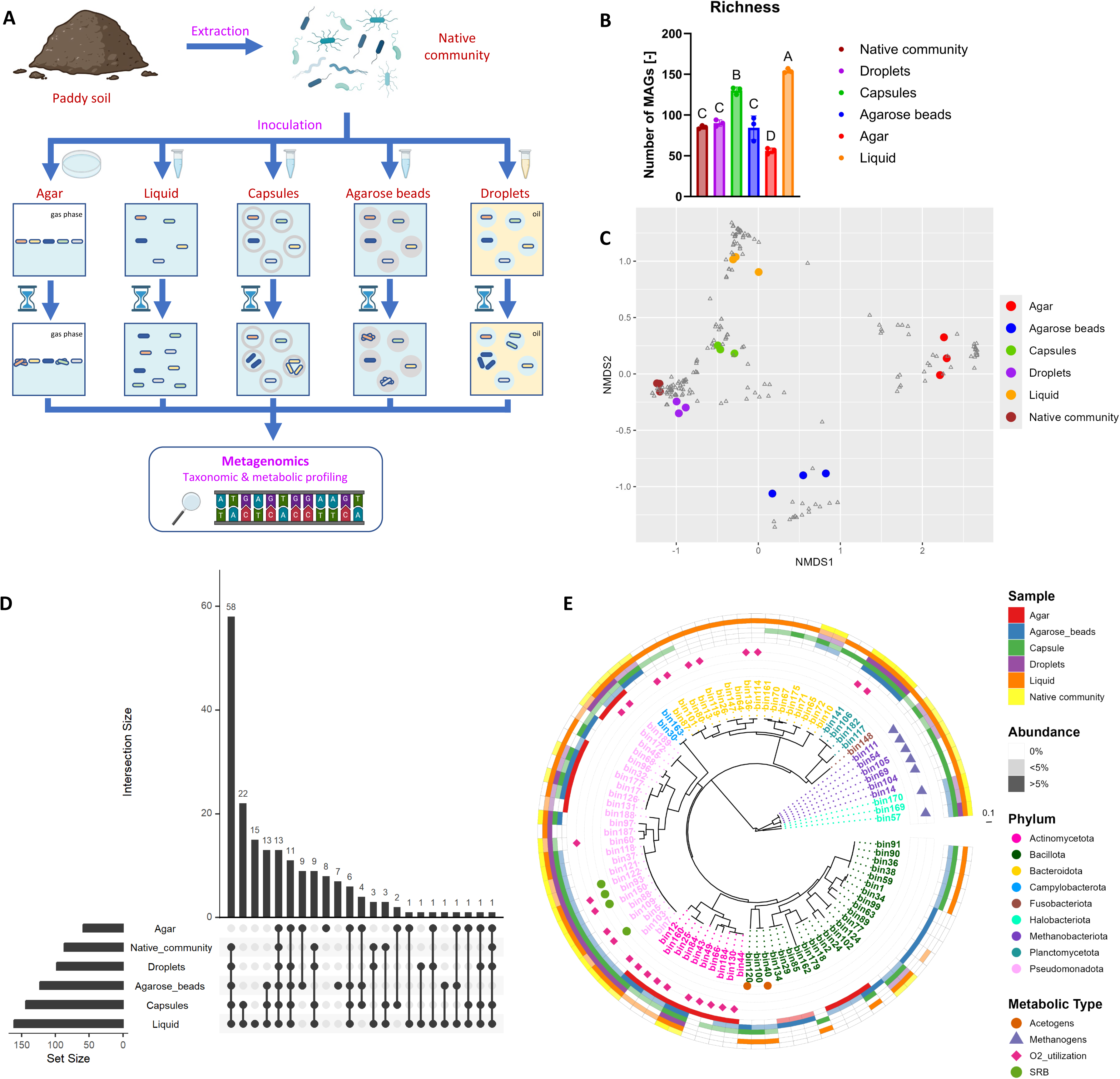
Comparison of different platforms to cultivate soil microbial taxa. (A) Schematic overview of the experiment. Microorganisms are extracted from a paddy soil (A_BH_) and grown anaerobically for 1 week in different cultivation platforms. After cultivation, microbial cells are harvested and all communities (initial and 1-week cultures) are profiled with metagenomics. (B) Richness of microbial communities. Bars lengths indicate mean richness values (number of MAGs) and error bars show standard deviation (N = 3 replicates). Individual replicate values are shown as dots. Different letters indicate significant difference at p < 0.05 between treatments (Tukey’s test). (C) Nonmetric multidimensional scaling (NMDS) plot illustrating differences across microbial communities, based on Bray-Curtis dissimilarity values derived from relative abundance data. The reads that did not map to any MAGs were removed from the analysis and relative abundance values were rescaled accordingly. Each MAG is shown as a triangle, whereas communities are depicted as colour dots. Different colours indicate different cultivation conditions (three replicates per condition). The stress value of the ordination was 0.0605. (D) UpSet plot showing taxonomic similarities across the different communities. The set size indicates, for each condition, the total number of MAGs who are present in at least one replicate. Intersection size shows the number of MAGs shared across the samples indicated with dots. (E) Circular maximum likelihood phylogenetic tree illustrating the diversity of microbial taxa (MAGs) present in the communities, based on the relative abundance values (average of 3 replicates). The colour of the MAG name indicates its taxonomy at the phylum level and its presence/absence in the different communities is shown in the outer circles (one specific colour is attributed to each sample type). Two abundance ranges (<5% and >5%) are specified with different colour shades. Symbols indicate MAGs with specific metabolic capabilities (acetogens, methanogens, aerobes and sulfate-reducers). The scale represents the branch length. SRB: sulfate-reducing bacteria.

### Soil microbial community undergoes temporal shifts during in-capsule cultivation

We cultivated soil microorganisms anaerobically within capsules and profiled the taxonomy over time. Gammaproteobacteria taxa grew rapidly and dominated the community within 24 h, but then decreased in relative abundance as other taxa (e.g. Actinomycetia, Clostridia, Bacteroidia) grew substantially between 24 h and 72 h (Fig. 3A). A closer look at the facultative anaerobic taxa (MAGs harbouring genes involved in oxygen respiration) showed a sharp increase in their relative abundance in the first 48 h, followed by a decrease, especially between 72 h and 144 h (Fig. 3B). The growth of strictly anaerobic taxa (Methanobacteria, Methanosarcinia and Desulfovibrionia) occurred during this time frame (Fig. 3A). In a parallel experiment where the spent medium was replaced with fresh medium after every time point, a similar result was observed (Fig. S11). We also attempted to further enrich slow-growing taxa by conducting similar experiments with longer incubation periods, but did not observe their growth (Fig. S12-13). Therefore, individual taxa exhibit distinct in-capsule growth dynamics.

**Figure 3:**
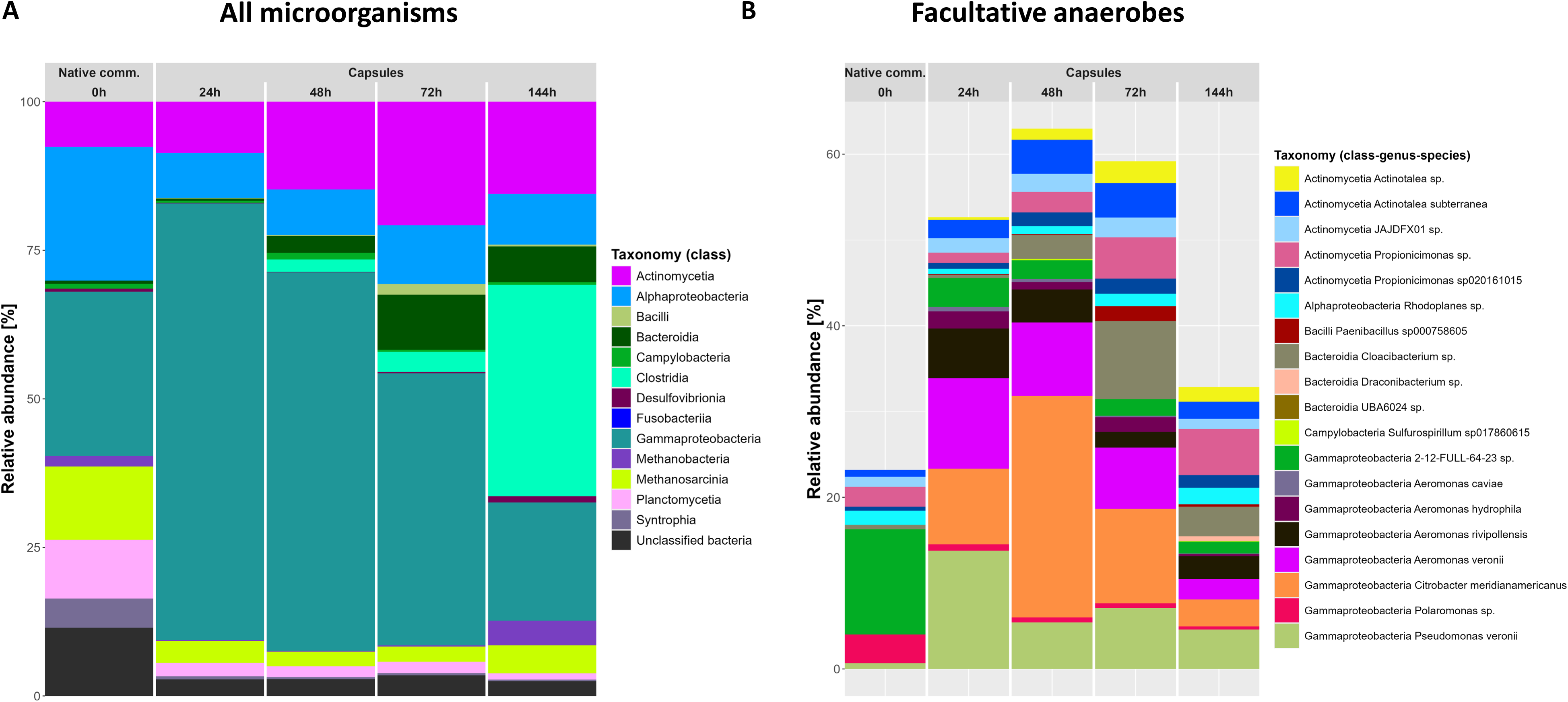
Dynamics of soil-derived microbial community within capsules over cultivation time. (A) Taxonomy barplot showing changes in the microbial community extracted from soil throughout cultivation in capsules in MSM10 medium. The MAGs are shown at the class level (see colour code in legend). The initial community, extracted from A_BH_ soil, is designated as ‘Extract’. Taxa labelled as ‘Unclassified bacteria’ are MAGs that were not assigned to a known phylum. (B) Taxonomy barplot showing changes in the abundance of facultative anaerobic taxa throughout cultivation in capsules. The taxonomy (family, genus and species) of each MAG is indicated, see colour code in legend.

### High-throughput capsule sorting enables the isolation of soil anaerobes

Similar to agarose beads and double-emulsion systems [26,32], we found that capsules can be sorted with FACS. Thus, we set out to isolate anaerobes using this high-throughput technique. Capsule-entrapped soil microorganisms were cultivated, the biomass fluorescently labeled and sorted into microwells to establish pure cultures. With this approach, we were able to isolate various soil-derived microorganisms, including the obligate anaerobe *Clostridium* J sp. 902363375 (Table 2).

**Table 2:** List of slow-growing anaerobes isolated from soil using capsules and FACS.

| Soil | Organisms in capsules | Medium (cultivation within capsules)<sup>a</sup> | Relative abundance within capsules at the time of sorting<sup>b</sup> | Medium (cultivation in microwell plate)<sup>a</sup> | Taxonomy of isolate<sup>c</sup> | Yield<sup>d</sup> |
| --- | --- | --- | --- | --- | --- | --- |
| A<sub>AH</sub> | Freshly extracted from soil | MSM1 | Not measured | MSM10 | *Clostridium* J sp. 902363375 | 1/25 (4.0%) |
| A<sub>BH</sub> | Freshly extracted from soil | DSMZ 311 medium, H<sub>2</sub>/CO<sub>2</sub> (80/20 v/v, 0.5 bar) | 0% | DSMZ 311 medium | *Terrisporobacter glycolicus* | 1/68 (1.5%) |
| A<sub>BH</sub> | Freshly extracted from soil | DSMZ 135 medium, H<sub>2</sub>/CO<sub>2</sub> (80/20 v/v, 0.5 bar) | 8.4% | DSMZ 135 medium | *Acetoanaerobium sticklandii* | 1/35 (2.9%) |
| A<sub>BH</sub> | Freshly extracted from soil | DSMZ 311 medium, H<sub>2</sub>/CO<sub>2</sub> (80/20 v/v, 0.5 bar) | Not measured | DSMZ 311 medium | *Paraclostridium* sp. | 1/24 (4.2%) |
| B<sub>BH</sub> | Enrichment culture (SRB) | Modified Postgate B medium, N<sub>2</sub>/CO<sub>2</sub> (20/80 v/v, 0.5 bar) | Not measured | Modified Postgate B medium, H<sub>2</sub>/CO<sub>2</sub> (20/80 v/v, 0.1 bar) | *Desulfovibrio desulfuricans* | 1/151 (0.7%) |
| B<sub>BH</sub> | Enrichment culture (SRB) | Modified Postgate B medium, N<sub>2</sub>/CO<sub>2</sub> (20/80 v/v, 0.5 bar) | Not measured | Modified Postgate B medium, H<sub>2</sub>/CO<sub>2</sub> (20/80 v/v, 0.1 bar) | *Lacrimispora xylanolytica* | 1/151 (0.7%) |
| B<sub>BH</sub> | Enrichment culture (methanogens) | Modified DSMZ 141c medium, H<sub>2</sub>/CO<sub>2</sub> (80/20 v/v, 2 bar) | Not measured | DSMZ 141c medium, H<sub>2</sub>/CO<sub>2</sub> (80/20 v/v, 0.1 bar) | *Nitratidesulfovibrio vulgaris* | 2/112 (1.8%) |

<sup>a</sup> Cultivation was performed at 30°C. When no gases are specified, the cultures were incubated in an anoxic chamber (97% N<sub>2</sub>, 3% H<sub>2</sub>). More information about the media can be found in SI.
<sup>b</sup> An aliquot of the capsules was stored for DNA extraction 3 h before FACS. Relative abundance values were obtained after mOTUs taxonomic assignment.
<sup>c</sup> The taxonomy of each isolate was determined by comparing its near-full length 16S rRNA gene (PCR- amplified with the 27F-1492R primers) to the GTDB database (release 202).
<sup>d</sup> Ratio between the number of isolates of the corresponding taxon and the total numbers of isolates obtained under the same conditions.

However, because most of the isolates obtained with the MSM medium were fast-growing, facultative anaerobic taxa (Fig. S14A), we repeated the experiment by using media containing reducing agents to promote slower-growing taxa. This approach proved successful, as we observed an enrichment of strict anaerobes within capsules (Fig. 4, Fig. S15), and could isolate some of them (*Terrisporobacter glycolicus*, *Acetoanaerobium sticklandii*, *Paraclostridium* sp.) (Table 2, Fig. S14B-D).

**Figure 4:**
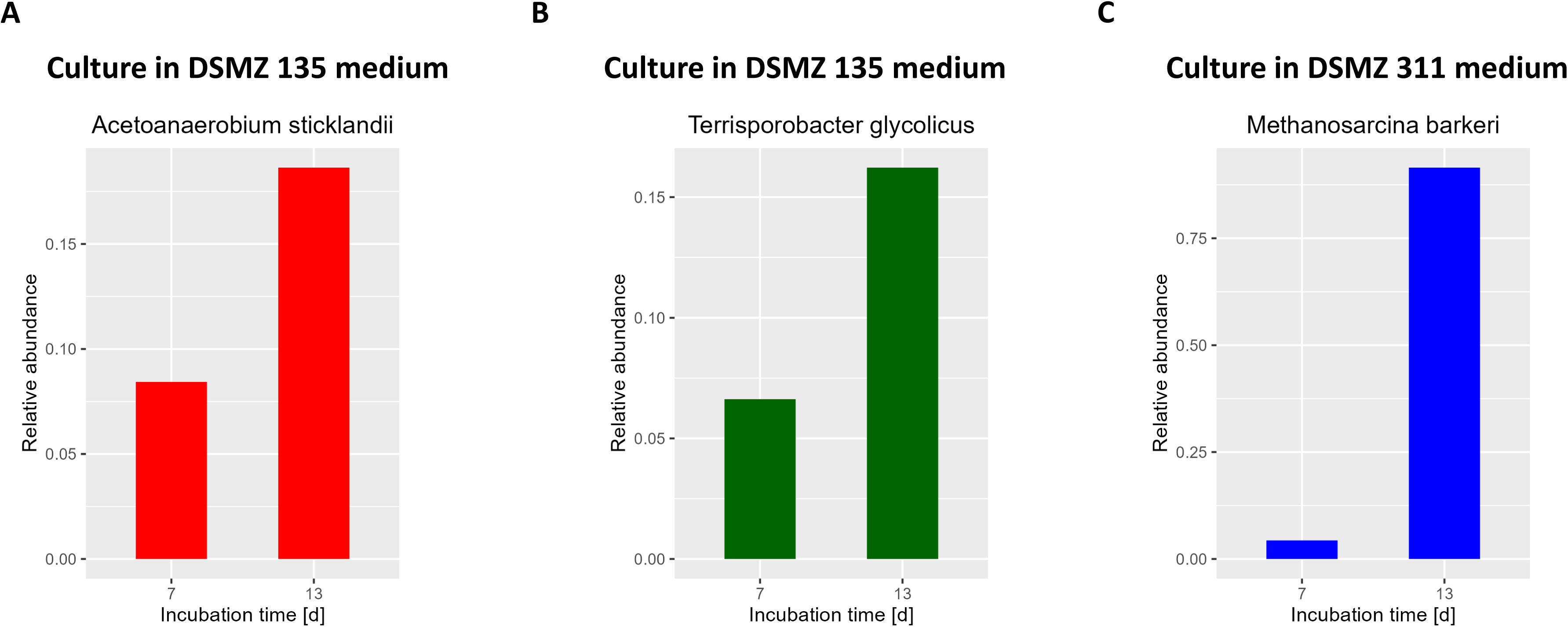
Growth of strictly anaerobic taxa within capsules in enrichment medium. Soil (A_BH_) microorganisms were individually entrapped in capsules and cultivated at 30°C in pre-reduced enrichment media (either DSMZ 135 or DSMZ 311 medium) in serum bottles filled with 80% H_2_, 20% CO_2_. After 7 and 13 days, an aliquot of the culture was sacrificed for taxonomic profiling (mOTUs). Barplots show the enrichment of (A) *Acetoanerobium sticklandii*, (B) *Terrisporobacter glycolicus* and (C) *Methanosarcina barkeri* within capsules, between 7 and 13 days of incubation. The data in (A) and (B) are from the same culture.

We further aimed to isolate SRB and methanogens, so we specifically enriched these microbial groups prior to encapsulation by incubating the soil under the appropriate selective conditions. This approach allowed us to isolate two SRB from soil (*Desulfovibrio desulfuricans* and *Nitratidesulfovibrio vulgaris*) (Table 2, Fig. S14E-F).

No methanogens were isolated from soil, although they were sometimes found in high abundance in capsules (Fig. 4C, Fig. S15). To assess the potential of our approach to isolate methanogens, we performed a control experiment where *Methanococcus maripaludis* was grown within capsules which were individually sorted in microwell plates. After a one-week incubation period, many wells showed growth of *M. maripaludis* (Fig. S16), while control wells (containing only medium) remained sterile. Thus, this strategy holds great potential to isolate diverse environmental anaerobes.

## Discussion

In this study, we presented a novel cultivation approach consisting of trapping and growing single microorganisms within capsules. We demonstrated the growth of phylogenetically and metabolically diverse anaerobes within capsules, including SRB, acetogens, and methanogens (Fig. 1C, 2E, 3A). Capsules act as microenvironments where microorganisms are physically separated but chemically interconnected. The selectively permeable shell of the capsule allows diffusion of nutrients and metabolites (including dissolved gases), enabling interspecies interactions (e.g. cross-feeding, competition). This likely explains why soil-derived cultures grown within capsules resemble batch liquid cultures (Fig. 2C-E).

However, capsules are compartments with limited space, restricting microbial growth to their maximum holding capacity. The decline in the growth of Gammaproteobacteria taxa following an early expansion phase may reflect the saturation of this available space (Fig. 3A-B), especially as medium replenishment did not allow further growth of these taxa (Fig. S11). Over a certain load, cell division may be inhibited by compressive forces or by a harmful accumulation of waste products. Therefore, in contrast with batch cultivation, capsules may advantageously prevent the proliferation of fast-growing microorganisms.

### Capsules enable the growth of diverse soil microbial taxa

Unlike agarose beads, which hold cells within a solid matrix, capsules allow microorganisms to grow as free-floating cells within a liquid core (Fig. S5). This distinction may explain the large differences between the cultures in these microcompartments (Fig. 2C-E, Fig. S9-10). Certain taxa such as Fusobacteriaceae gen. sp. (bin148), *Anaerorhabdus* sp. (bin 162) and many members in the phylum Bacteroidota, were found to grow within capsules and in batch liquid cultures but not within agarose beads (Fig. 2E, Table S2). Microbial cultivation within agarose beads was strongly biased towards specific MAGs, in particular members of the classes Clostridia and Bacilli (Fig. S10-11). This may indicate that spore-forming bacteria are better fitted to thrive within gel matrices. In addition, nine taxa grew exclusively on agar and within agarose beads (Fig. 2D), which suggests surface attachment improved their fitness. Such mechanisms have been observed in biofilm-forming bacteria through the upregulation of genes involved in stress response and virulence [34,35]. Alternatively, these microorganisms may be able to enzymatically degrade agarose (also a major component of agar) and utilize it as source of carbon or energy [36].

Consistent with the Great Plate Count Anomaly, only a small fraction of soil microorganisms grew on agar (Fig. 2B). In particular, no archaea and no member of the phyla Bacteroidota, Planctomycetota and Fusobacteriota were found in the agar cultures, whereas Actinomycetota members were highly abundant (Fig. 2E). This bias toward facultative anaerobes may result from the presence of trace levels of oxygen at the gas-liquid interface or from the formation of hydrogen peroxide upon autoclaving phosphate and agar together [12,37,38].

Recently, McCully *et al.* stressed the importance of nutrient privatization for cultivating slow-growing taxa, some of which (Methanobacteria, Negativicutes) they enriched from human stool using a double-emulsion platform (water-in-oil-in-water) [26]. However, the authors also proposed that chemical diffusion may occur to a limited extent through the thin oil layer and allow the growth of certain microorganisms via cross-feeding. Here, although similar taxa (Methanobacteria, Methanosarcinia, Desulfovibrionia and Negativicutes) were observed within capsules and in batch liquid culture, where nutrient sharing and interspecies interactions (e.g. competition) take place, we have not evidenced their enrichment in single emulsion (droplets) (Fig. 2E, 3A, S9-10). A greater nutrient privatization is expected in droplets as compared to double emulsion, because of the significantly thicker oil barrier separating aqueous compartments (i.e. chemical diffusion is slower). The preclusion of interspecies interactions may explain why some organisms, such as the hydrogenotrophic methanogen *Methanobacterium lacus* (bin69, bin104, and bin14), did not grow within droplets (Fig. 2E). Therefore, we propose that, under certain conditions, interspecies interactions are more important than nutrient privatization for enriching slow-growing taxa.

Positive interspecies interactions may have enabled the growth of certain taxa within capsules. For instance, we evidenced the growth of a novel species in the Negativicutes genus *Sporomusa* (bin120) (Fig. 2E, Table S2), whose members can perform acetogenesis and fermentation of *N*-methyl compounds [39]. Metagenomic data indeed reveal the presence of genes involved in the Wood-Ljungdahl pathway in this organism, thus requiring CO_2_ and H_2_ (Table S2) These growth substrates are likely produced by other organisms, typically fermenters, before being utilized by *Sporomusa* sp.. We also evidenced the growth of methanogens harbouring genes for all known methanogenic pathways (Fig. 2E, Table S2). Methanogenic substrates (e.g. acetate, H_2_) may similarly be released by fermentative organisms, which would explain why the growth of methanogens occurs only after an increase in abundance of Bacteroidia and Clostridia taxa (Fig. 3A, S11). In other experiments, the lack of sufficient fermentative activity may be responsible for the inability of SRB and methanogens to grow (Fig. S12-13). Finally, facultative anaerobes may set the stage for the growth of strict anaerobes by consuming residual oxygen, lowering the redox potential.

### Novel taxa grow in capsules

This study also revealed the presence of yet-uncultured but environmentally relevant taxa in soil-derived cultures. For instance, peculiar Burkholderiales bacteria (bin93, bin150 and bin5) were found with the potential to fully reduce sulfate to sulfide while also being capable of respiring oxygen (Fig. 2E, Table S2). Similar taxa had been reported in sulfate-rich zones within estuary sediments [40]; here, they may possibly occupy sulfate-rich soil microsites. Also, we identified microbial taxa in the candidate genus *Anammoximicrobium* (bin141, bin106 and bin182) (Fig. 2E, Table S2), i.e. putative anaerobic ammonium oxidizers, which had previously been observed in wastewater [41]. While these taxa were all present in capsule-grown cultures, none were detected in agar-grown cultures.

### Isolation of anaerobes using capsules

Capsules can be sorted individually to isolate microorganisms, including obligate anaerobes, in a high-throughput manner. This is of high interest because anaerobes are traditionally isolated with labour-intensive techniques (e.g. dilution-to-extinction, Hungate roll-tube). One advantage of FACS is that only a very low biomass (< 100 cells) is required for sorting. In comparison, conventional isolation methods based on visual screening of colonies may require millions of cells. We also found that, like agarose beads [32], capsules break open immediately after sorting, which conveniently removes the need for dissolving them before growing isolates in liquid medium.

The comparison across platforms showed that both capsules and droplets can effectively recapitulate the initial soil community (Fig. 2B-E). However, droplets cannot be readily separated from the oil phase (due to the absence of a rigid shell) and therefore, cannot be FACS sorted. Double-emulsion droplets (McCully et al., 2023) can be sorted but still suffer from the requirement to add the DNA stain upon encapsulation as opposed to after growth as for capsules.

Here, we successfully isolated soil anaerobes via capsule sorting (Table 2). The incubation of capsules in enrichment media prior to sorting allowed us to enrich (Fig. 4, Fig. S15) and isolate (Table 2) slow-growing microorganisms, including two putative acetogens, *Acetoanaerobium sticklandii* and *Terrisporobacter glycolicus* [42]. We also enriched for SRB and methanogens before encapsulation, which allowed us to isolate *D. desulfuricans* and *N. vulgaris*, two SRB. However, no methanogens were isolated, despite their ability to grow within capsules (Fig. 4C). Some methanogens are known to initiate growth only when the redox potential is sufficiently low (< 0 mV) [43,44]. Since FACS was not performed under anoxic conditions, we suspect that this short exposure to air may have inhibited their growth. Yet, an isolation workflow conducted under similar conditions with the methanogen *M. maripaludis* resulted in its growth in the microwell plate (Fig. S16). Thus, soil methanogens may require specific growth factors released by other microorganisms which were not provided in our medium in the microplate.

In previous studies, microbial growth had been shown to occur within capsules but only with readily cultivable model organisms under aerobic conditions [27,45,46]. This work demonstrates that this platform can further be used to study unknown environmental taxa. For instance, single-cell compartmentalization, cultivation within capsules and shell dissolution steps may be repeatedly performed to enrich rare taxa. Several microorganisms may also be co-encapsulated to enrich elusive syntrophic taxa.

Capsules had, to our knowledge, not previously been used in combination with FACS to isolate microorganisms. Here, we demonstrated the strong potential of this approach by isolating slow-growing anaerobic bacteria from soil. In contrast, agarose beads and droplets have been employed for isolation, but we show that only a small subset of microbial taxa grow in these systems as compared to capsules.

We envision that *in situ* incubation of capsules prior to sorting could facilitate the isolation of novel taxa. Ben-Dov *et al.* have previously shown the potential of *in situ* cultivation by reporting the growth of novel coral-associated microorganisms within polysulfone-coated agar beads, although this has not resulted in isolation [47]. Here, capsules could be incubated in diffusion chambers directly placed into the native environment [48,49]. The success of the isolation would depend on the ability of the organisms to grow in the microwell plate after FACS, which could be aided by the use of dedicated environment-mimicking media. Furthermore, capsules may facilitate the isolation of hyperthermophilic microorganisms for which agar, melting at 85°C, cannot be used [50]. Here we have shown the growth of a thermophile (*Thermoanaerobacter kivui*) at 60°C (Fig. 1C), but capsules can even stand heat up to 100°C (Atrandi Biosciences, pers. commun., 8 November 2024).

## Acknowledgments

We acknowledge Natalia Jakus for assistance with soil sampling, Wenyu Gu for providing strain MM901, Yolanda Schaerli and Hannes Peter for insightful discussions, and the EPFL PTCF for help with FACS.

## Data Availability

All sequencing data are accessible under the NCBI BioProject PRJNA1190322. The code used in this study is available on <u>GitHub</u>.

---

Figure S1: **Graphical summary of the study.**

Figure S2: **Experimental setup used to produce capsules under anoxic conditions**. Left: Capsules are generated with an Onyx microfluidic platform (Atrandi Biosciences) which includes syringe pumps and a microscope. Core and shell reagents (along with the microbial cells) are injected in a microfluidic chip, and the resulting emulsion (which will result in capsules) is collected in an microtube. Right: Overall view of the setup, where capsules are produced inside an anoxic chamber with the Onyx platform, while images and flow rates are monitored on an external device connected through Wi-Fi.

Figure S3: **FACS scatter plots with gates on microcolony-containing capsules.** A first gate (P1) was defined to exclude smaller particles (e.g. free-floating cells or debris), whereas the second gate (P2) further selects the capsules with high SYTO-9 fluorescence signal prior to sorting. Left: analysis of capsules with no microbial growth (incubation in PBS). Right: analysis of capsules with microbial growth (incubation in medium).

Figure S4: **Description of different types of microcompartments**.

Figure S5: **Microscopy images of microorganisms trapped in capsules.** *D. vulgaris* and *E*. *coli* TB205 were imaged immediately after encapsulation and emulsion breaking (no incubation) whereas *Shewanella oneidensis* MR-1 was imaged after a 24h incubation period in M9 medium at 30°C (see Table 1 for details). For each microbial strain, brightfield (left) and fluorescent (right) images were captured with a microscope (Eclipse Ni-E, Nikon) and disposed side by side. The microbial cells were stained by incubating the sample with SYTO-9 (1 µM) for 10 min in the dark, except for *E*. *coli* TB205 which constitutively expresses a gene to produce the fluorescent protein mCherry, and thus did not require staining.

Figure S6: **Taxonomic composition of the microbial community in soil AAH.** The taxonomy barplot shows the relative abundance values of indicated microbial taxa at the phylum level. Taxonomy assignment was conducted based on the full-length 16S rRNA genes using the SILVA database.

Figure S7: **Growth of soil microorganisms within microcompartments in a soil-derived minimal medium.** After extraction from soil, single cells were trapped in w-o droplets, capsules and agarose beads and incubated at 30°C in MSM10. Images taken after different times reveal the growth of organisms in droplets (left), capsules (middle) and agarose beads (right). Arrows in the left figure point to microbial cells.

Figure S8: **Nonmetric multidimensional scaling (NMDS) plot illustrating differences across microbial communities, based on Bray-Curtis dissimilarity values derived from relative abundance data (mOTUs).** mOTUs reads which did not map to any known reference were removed from the analysis and relative abundance values were rescaled accordingly. Each OTU is shown as a triangle, whereas communities are depicted as colours dots. Different colours indicate different cultivation conditions (three replicates per condition). The stress value of the ordination was 0.0589.

Figure S9: **Taxonomy barplot of the soil microbial communities across different cultivation platforms (MAGs).** Relative abundance values are shown as means of three replicates. Taxonomy is displayed at the class level. The native (initial) community is labelled as ‘Extract’. MAGs with relative abundance >3% are displayed. Reads which do not map to any MAG are not shown. The ‘Unclassified’ label indicates MAGs which map to bacteria of unknown phyla.

Figure S10: **Taxonomy barplot of the soil microbial communities across different cultivation platforms (mOTUs).** Relative abundance values are shown as means of three replicates. Taxonomy is displayed at the class level. The native (initial) community is labelled as ‘Extract’. OTUs with relative abundance >1% are displayed. The ‘Unassigned’ label indicates OTUs which do not map to any reference in the mOTUs database.

Figure S11: **Temporal dynamics of microbial communities growing within capsules in medium replenished after each sampling.** Taxonomy barplot showing changes in the microbial community throughout cultivation. After each time point, the old medium was replaced with fresh medium. The MAGs are shown at the class level (see colour code in legend). The initial community, extracted from A_BH_ soil, is designated as ‘Extract’. Taxa labelled as ‘Unclassified bacteria’ are MAGs that were not assigned to a known phylum.

Figure S12: **Temporal dynamics of microbial communities growing within capsules (soil A_BH_, MSM1 medium) over a longer time period.** Taxonomy barplot showing changes in the microbial community throughout cultivation. Relative abundance values are shown as means of three replicates. MAGs are shown at the phylum level (see colour code in legend). The initial community is designated as ‘Extract’. Taxa labelled as ‘Unclassified bacteria’ are MAGs that were not assigned to a known phylum.

Figure S13: **Temporal dynamics of microbial communities growing within capsules (soil B_BH_, MSM1 medium) over a longer time period.** Taxonomy barplot showing changes in the microbial community throughout cultivation. Relative abundance values are shown as means of three replicates. MAGs are shown at the phylum level (see colour code in legend). The initial community is designated as ‘Extract’. Taxa labelled as ‘Unclassified bacteria’ are MAGs that were not assigned to a known phylum.

Figure S14: **Pie charts of all isolates obtained across experiments listed in Table 2**. Each chart corresponds to a separate experiment. The isolates listed in Table 2 are highlighted as exploded slices, with genus-level taxonomy displayed.

Figure S15: **Taxonomy barplots showing the changes in relative abundance of capsule-grown microorganisms (derived from soil A_BH_) after 7 and 13 days of incubation in different media.** The taxonomy was profiled with mOTUs.

Figure S16: **Microscopy image of *Methanococcus maripaludis* MM901 cells growing in the microwell plate after FACS.** Single-cell encapsulation was performed on a pure culture of *Methanococcus maripaludis* MM901. The capsules were incubated in DSMZ 141c medium for 1 month after which single capsules were sorted and distributed in a 96-well plate filled with DSMZ 141c medium. Immediately after sorting, the 96-well plate was incubated in an anaerobic jar filled with H_2_/CO_2_ (80/20 v/v) headspace. After one week of incubation, the plate was taken out of the jar, and growth could be seen visually in several wells. The image shown was taken from one of these wells.

Figure S17: **Goodness of fit and Shepard plots for the NMDS.** Left: the goodness of fit of the representation is illustrated for each sample by the diameter of the circles around them (larger circles indicating poorer fits) whereas red crosses indicate MAGs. Right: the Shepard plot shows the agreement (high R-squared values) between the original distance matrix and the ordination representation.

Figure S18: **Analysis of similarities (ANOSIM) plot showing dissimilarity between and within communities in the NMDS ordination.** The bold line within each box represents the median. The lower edge of the box marks the 25^th^ percentile, while the upper edge marks the 75^th^ percentile. Whiskers extend to the most extreme data points within 1.5 times the interquartile range from the box.

Table S2: **Summary of taxonomic and metabolic profiles of metagenome-assembled genomes in soil-derived cultures**.

## References

1. Lewis WH, Tahon G, Geesink P et al. Innovations to culturing the uncultured microbial majority. Nat Rev Microbiol 2021;19:225–40.

2. BacDive (DSMZ) dashboard, https://bacdive.dsmz.de/dashboard (9 November 2024, date last accessed).

3. Louca S, Mazel F, Doebeli M et al. A census-based estimate of earth’s bacterial and archaeal diversity. PLoS Biol 2019;17:e3000106.

4. LPSN (DSMZ) numbers, https://lpsn.dsmz.de/text/numbers (9 November 2024, date last accessed).

5. Schloss PD, Girard RA, Martin T et al. Status of the archaeal and bacterial census: An update. Delong EF, McFall-Ngai MJ (eds.). MBio 2016;7, DOI: 10.1128/mBio.00201-16.

6. Rinke C, Schwientek P, Sczyrba A et al. Insights into the phylogeny and coding potential of microbial dark matter. Nature 2013;499:431–7.

7. Lloyd KG, Steen AD, Ladau J et al. Phylogenetically Novel Uncultured Microbial Cells Dominate Earth Microbiomes. mSystems 2018;3, DOI: 10.1128/msystems.00055-18.

8. Staley JT, Konopka A. Measurement of in situ activities of nonphotosynthetic microorganisms in aquatic and terrestrial habitats. Annu Rev Microbiol 1985;39:321–46.

9. Connon SA, Giovannoni SJ. High-throughput methods for culturing microorganisms in very-low-nutrient media yield diverse new marine isolates. Appl Environ Microbiol 2002;68:3878–85.

10. Bruns A, Cypionka H, Overmann J. Cyclic AMP and acyl homoserine lactones increase the cultivation efficiency of heterotrophic bacteria from the central Baltic Sea. Appl Environ Microbiol 2002;68:3978–87.

11. Janssen PH, Yates PS, Grinton BE et al. Improved culturability of soil bacteria and isolation in pure culture of novel members of the divisions Acidobacteria, Actinobacteria, Proteobacteria, and Verrucomicrobia. Appl Environ Microbiol 2002;68:2391–6.

12. Tanaka T, Kawasaki K, Daimon S et al. A hidden pitfall in the preparation of agar media undermines microorganism cultivability. Appl Environ Microbiol 2014;80:7659–66.

13. Huang Y, Sheth RU, Zhao S et al. High-throughput microbial culturomics using automation and machine learning. Nat Biotechnol 2023;41:1424–33.

14. Lagier JC, Khelaifia S, Alou MT et al. Culture of previously uncultured members of the human gut microbiota by culturomics. Nat Microbiol 2016;1:1–8.

15. Li S, Lian WH, Han JR et al. Capturing the microbial dark matter in desert soils using culturomics-based metagenomics and high-resolution analysis. npj Biofilms Microbiomes 2023;9:1–14.

16. Hanišáková N, Vítězová M, Rittmann SKMR. The Historical Development of Cultivation Techniques for Methanogens and Other Strict Anaerobes and Their Application in Modern Microbiology. Microorganisms 2022;10:412.

17. Hungate RE. Chapter IV A Roll Tube Method for Cultivation of Strict Anaerobes. Methods Microbiol 1969;3:117–32.

18. In’t Zandt MH, De Jong AEE, Slomp CP et al. The hunt for the most-wanted chemolithoautotrophic spookmicrobes. FEMS Microbiol Ecol 2020;94:64.

19. Cho JC, Giovannoni SJ. Cultivation and Growth Characteristics of a Diverse Group of Oligotrophic Marine Gammaproteobacteria. Appl Environ Microbiol 2004;70:432–40.

20. Kim S, Park MS, Song J et al. High-throughput cultivation based on dilution-to-extinction with catalase supplementation and a case study of cultivating acI bacteria from Lake Soyang. J Microbiol 2020;58:893– 905.

21. Rappé MS, Connon SA, Vergin KL et al. Cultivation of the ubiquitous SAR11 marine bacterioplankton clade. Nat 2002 4186898 2002;418:630–3.

22. Ma L, Datta SS, Karymov MA et al. Individually addressable arrays of replica microbial cultures enabled by splitting SlipChips. Integr Biol (Camb) 2014;6:796–805.

23. Ma L, Kim J, Hatzenpichler R et al. Gene-targeted microfluidic cultivation validated by isolation of a gut bacterium listed in human microbiome project’s most wanted taxa. Proc Natl Acad Sci U S A 2014;111:9768–73.

24. Nichols D, Cahoon N, Trakhtenberg EM et al. Use of ichip for high-throughput in situ cultivation of “uncultivable microbial species▽. Appl Environ Microbiol 2010;76:2445–50.

25. Zengler K, Toledo G, Rappé M et al. Cultivating the uncultured. Proc Natl Acad Sci U S A 2002;99:15681–6.

26. McCully AL, Loop Yao M, Brower KK et al. Double emulsions as a high-throughput enrichment and isolation platform for slower-growing microbes. ISME Commun 2023 31 2023;3:1–9.

27. Leonaviciene G, Leonavicius K, Meskys R et al. Multi-step processing of single cells using semi-permeable capsules. Lab Chip 2020;20:4052–62.

28. Mastiani M, Firoozi N, Petrozzi N et al. Polymer-Salt Aqueous Two-Phase System (ATPS) Micro-Droplets for Cell Encapsulation. Sci Reports 2019 91 2019;9:1–9.

29. Baghbanbashi M, Shiran HS, Kakkar A et al. Recent advances in drug delivery applications of aqueous two-phase systems. PNAS Nexus 2024;3, DOI: 10.1093/PNASNEXUS/PGAE255.

30. Semi-Permeable Capsules (Atrandi Biosciences), https://atrandi.com/semi-permeable-capsules (20 November 2024, date last accessed).

31. Atrandi Biosciences. SPC Innovator Kit User Guide. Document No. DGPM02323062001, Version 5.

32. Schaerli Y. Bacterial Microcolonies in Gel Beads for High-throughput Screening. Bio-protocol 2018;8, DOI: 10.21769/BIOPROTOC.2911.

33. Schaerli Y. Bacterial Microcolonies in Gel Beads for High-throughput Screening. Bio-Protocol 2018;8, DOI: 10.21769/bioprotoc.2911.

34. Siryaporn A, Kuchma SL, O’Toole GA et al. Surface attachment induces Pseudomonas aeruginosa virulence. Proc Natl Acad Sci U S A 2014;111:16860–5.

35. Wang L, Wu Y, Cai P et al. The attachment process and physiological properties of Escherichia coli O157:H7 on quartz. BMC Microbiol 2020;20:355.

36. Kováč J, Kushkevych I. New modification of cultivation medium for isolation and growth of intestinal sulfate-reducing bacteria. 2017.

37. Kawasaki K, Kamagata Y. Phosphate-catalyzed hydrogen peroxide formation from agar, gellan, and κ-carrageenan and recovery of microbial cultivability via catalase and pyruvate. Appl Environ Microbiol 2017;83:e01366–17.

38. Watanabe M, Igarashi K, Kato S et al. Critical Effect of H2O2 in the Agar Plate on the Growth of Laboratory and Environmental Strains. Microbiol Spectr 2022;10, DOI: 10.1128/SPECTRUM.03336-22/SUPPL_FILE/SPECTRUM.03336-22-S0001.PDF.

39. Möller B, Oßmer R, Howard BH et al. Sporomusa, a new genus of gram-negative anaerobic bacteria including Sporomusa sphaeroides spec. nov. and Sporomusa ovata spec. nov. Arch Microbiol 1984;139:388–96.

40. Baker BJ, Lazar CS, Teske AP et al. Genomic resolution of linkages in carbon, nitrogen, and sulfur cycling among widespread estuary sediment bacteria. Microbiome 2015;3:14.

41. Khramenkov S V, Kozlov MN, Kevbrina M V et al. A novel bacterium carrying out anaerobic ammonium oxidation in a reactor for biological treatment of the filtrate of wastewater fermented sludge. Microbiology 2013;82:628–36.

42. Böer T, Schüler MA, Lüschen A et al. Isolation and characterization of novel acetogenic strains of the genera Terrisporobacter and Acetoanaerobium. Front Microbiol 2024;15, DOI: 10.3389/fmicb.2024.1426882.

43. Fetzer S, Conrad R. Effect of redox potential on methanogenesis by Methanosarcina barkeri. Arch Microbiol 1993;160:108–13.

44. Hirano S, Matsumoto N, Morita M et al. Electrochemical control of redox potential affects methanogenesis of the hydrogenotrophic methanogen Methanothermobacter thermautotrophicus. Lett Appl Microbiol 2013;56:315–21.

45. Steinacher M, Cont A, Du H et al. Monodisperse Selectively Permeable Hydrogel Capsules Made from Single Emulsion Drops. ACS Appl Mater Interfaces 2021;13:15601–9.

46. Di Girolamo S, Puorger C, Lipps G. Stable and selective permeable hydrogel microcapsules for high-throughput cell cultivation and enzymatic analysis. Microb Cell Fact 2020;19:1–13.

47. Ben-Dov E, Kramarsky-Winter E, Kushmaro A. An in situ method for cultivating microorganisms using a double encapsulation technique. FEMS Microbiol Ecol 2009;68:363–71.

48. Bollmann A, Lewis K, Epstein SS. Incubation of environmental samples in a diffusion chamber increases the diversity of recovered isolates. Appl Environ Microbiol 2007;73:6386–90.

49. Kaeberlein T, Lewis K, Epstein SS. Isolating “Uncultivable” Microorganisms in Pure Culture in a Simulated Natural Environment. Science (80- ) 2002;296:1127–9.

50. Das N, Triparthi N, Basu S et al. Progress in the development of gelling agents for improved culturability of microorganisms. Front Microbiol 2015;6:147147.

51. Viacava K, Qiao J, Janowczyk A et al. Meta-omics-aided isolation of an elusive anaerobic arsenic-methylating soil bacterium. ISME J 2022;16:1740–9.

52. Sambrook J, Russell DW. Molecular Cloning: A Laboratory Manual. Third Edition. United States Cold Spring Harb Lab Press 2001:4.35.

53. Bergmiller T, Andersson AMC, Tomasek K et al. Biased partitioning of the multidrug efflux pump AcrAB-TolC underlies long-lived phenotypic heterogeneity. Science (80- ) 2017;356:311–5.

54. Weghoff MC, Müller V. CO metabolism in the thermophilic acetogen Thermoanaerobacter kivui. Appl Environ Microbiol 2016;82:2312–9.

55. Costa KC, Wong PM, Wang T et al. Protein complexing in a methanogen suggests electron bifurcation and electron delivery from formate to heterodisulfide reductase. Proc Natl Acad Sci U S A 2010;107:11050–5.

