## Supplementary Figure 1 for "High-throughput cultivation and isolation of environmental anaerobes using selectively permeable hydrogel capsules"

### Comparison of different cultivation platforms (Fig. 2)

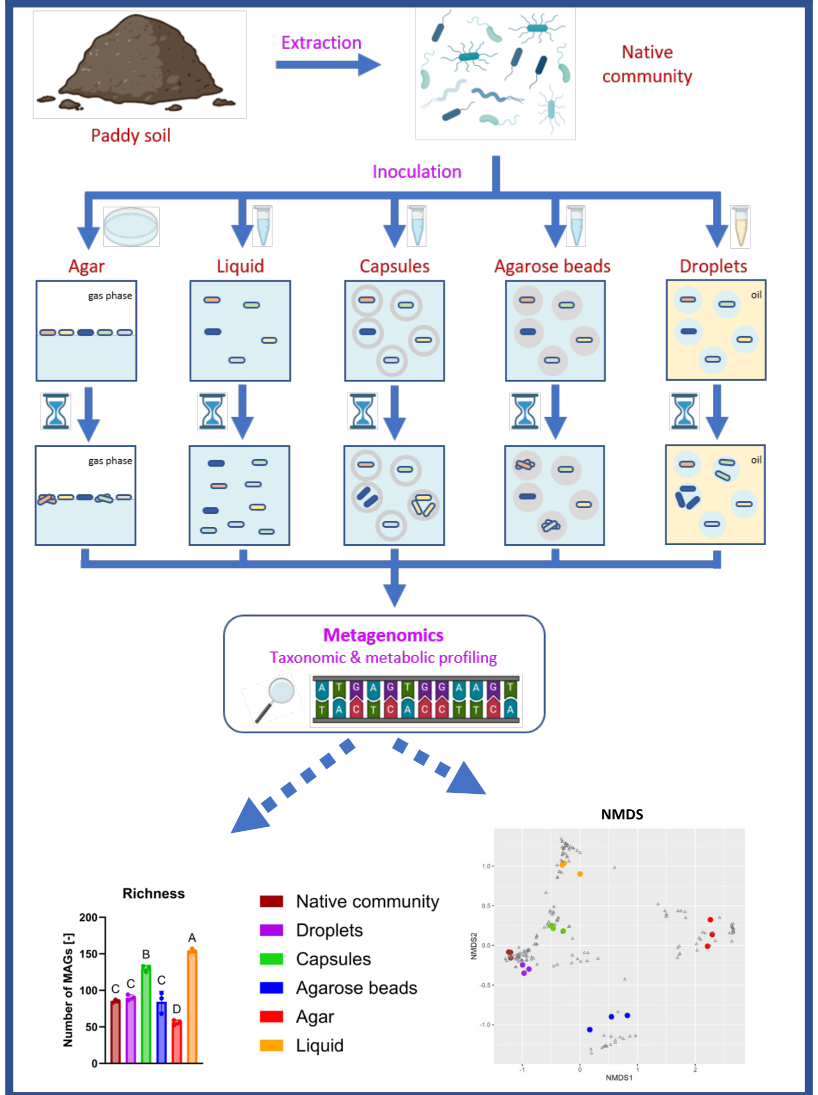


Soil microorganisms are grown in a minimal medium using different cultivation platforms. After incubation, the communities are compared.

### Changes in microbial community over time in capsules (Fig. 3)

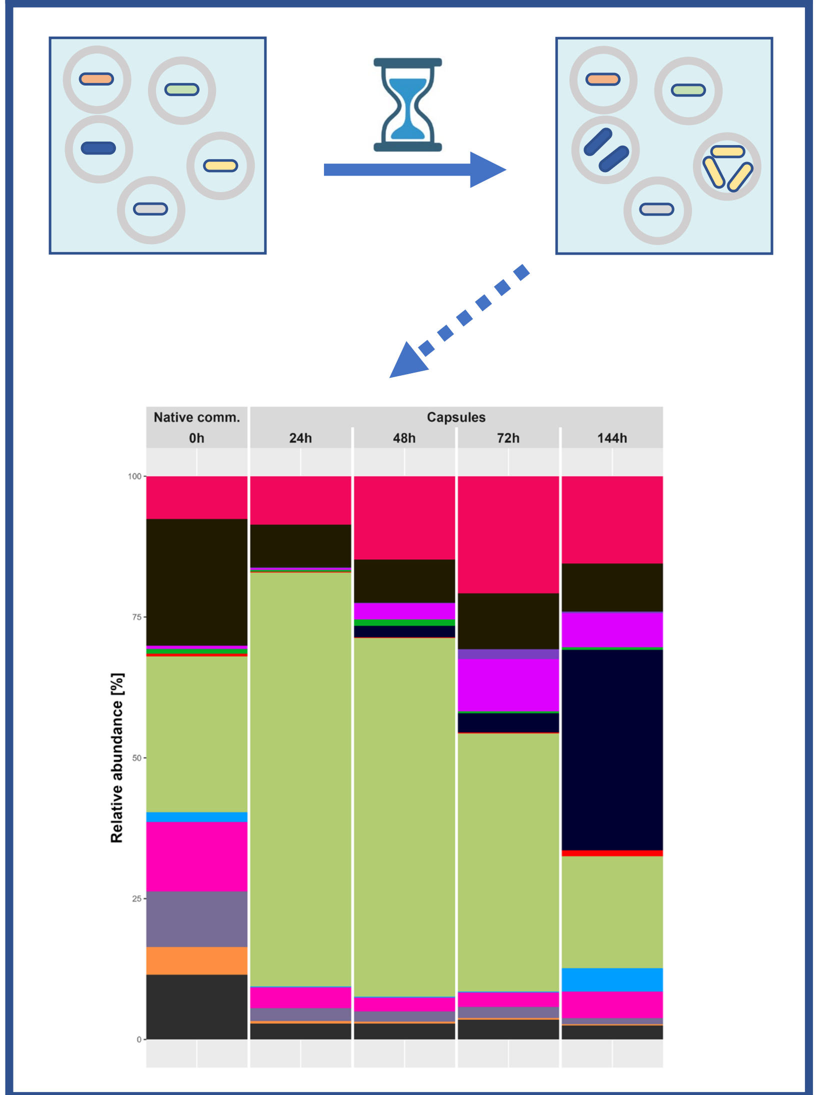


Soil microorganisms are grown in hydrogel capsules and the taxonomy is profiled after different incubation periods.

### Isolation of anaerobes (Table 2)

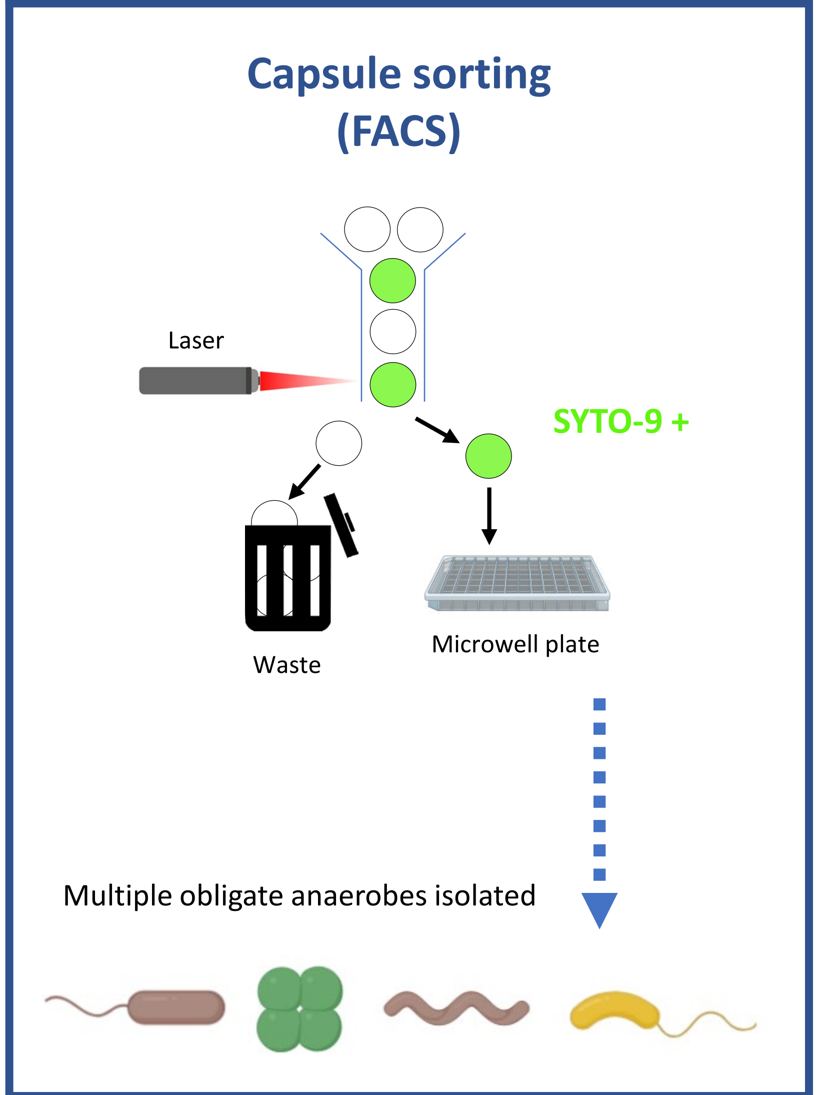


After encapsulation and growth, clonal populations can be isolated by sorting capsules (FACS) in microwell plates. The biomass is stained with SYTO-9.
