## Supplementary figures and images for "High-throughput cultivation and isolation of environmental anaerobes using selectively permeable hydrogel capsules"

### Supplementary Figure 2

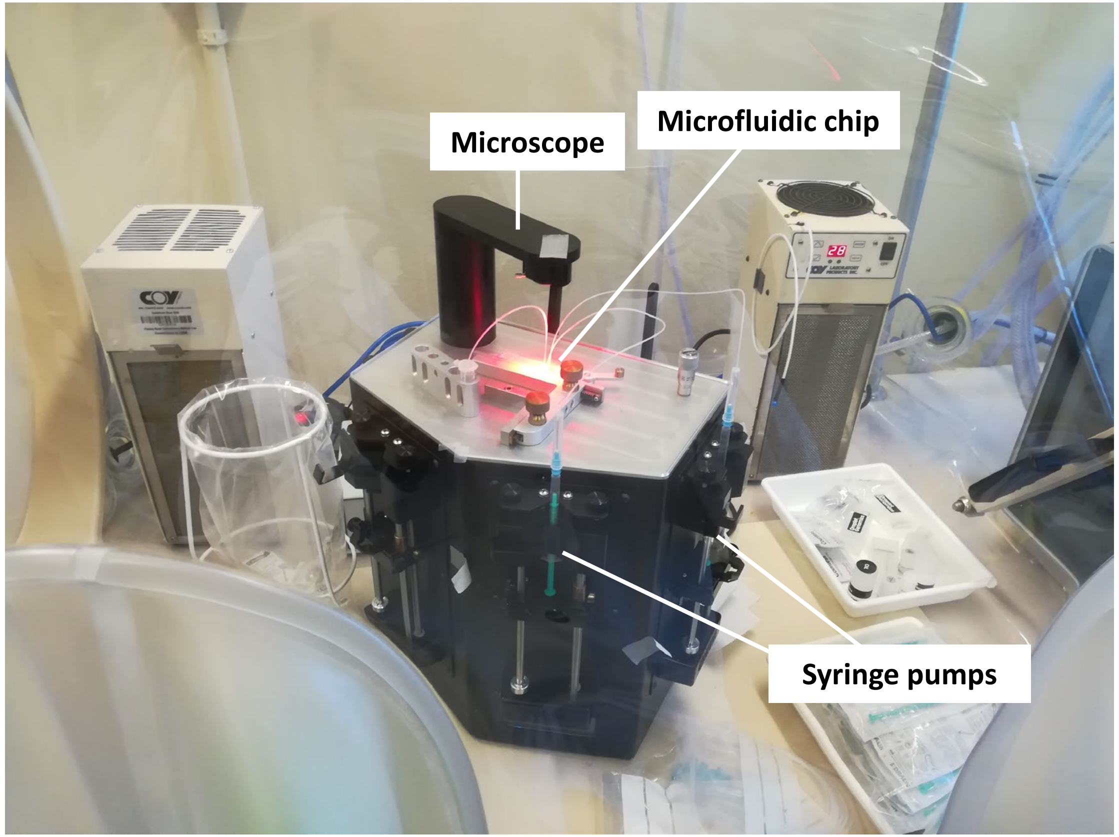

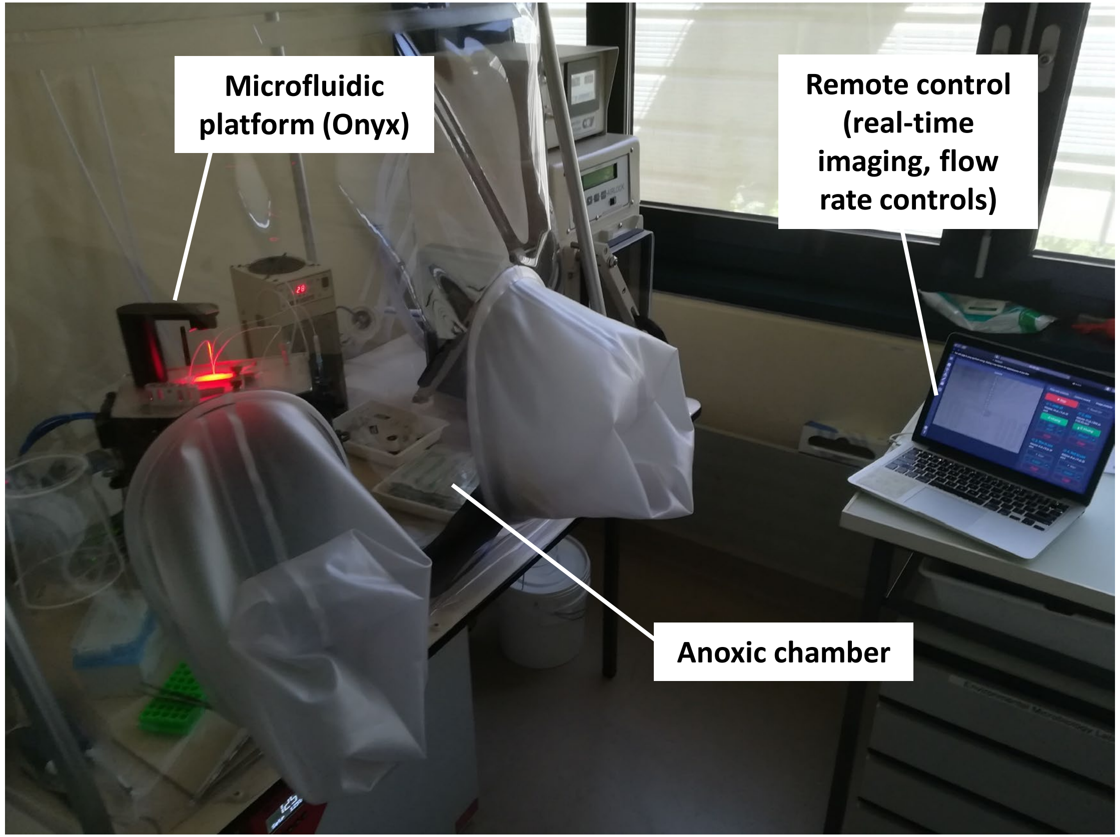

### Supplementary Figure 3

#### Capsules (PBS)

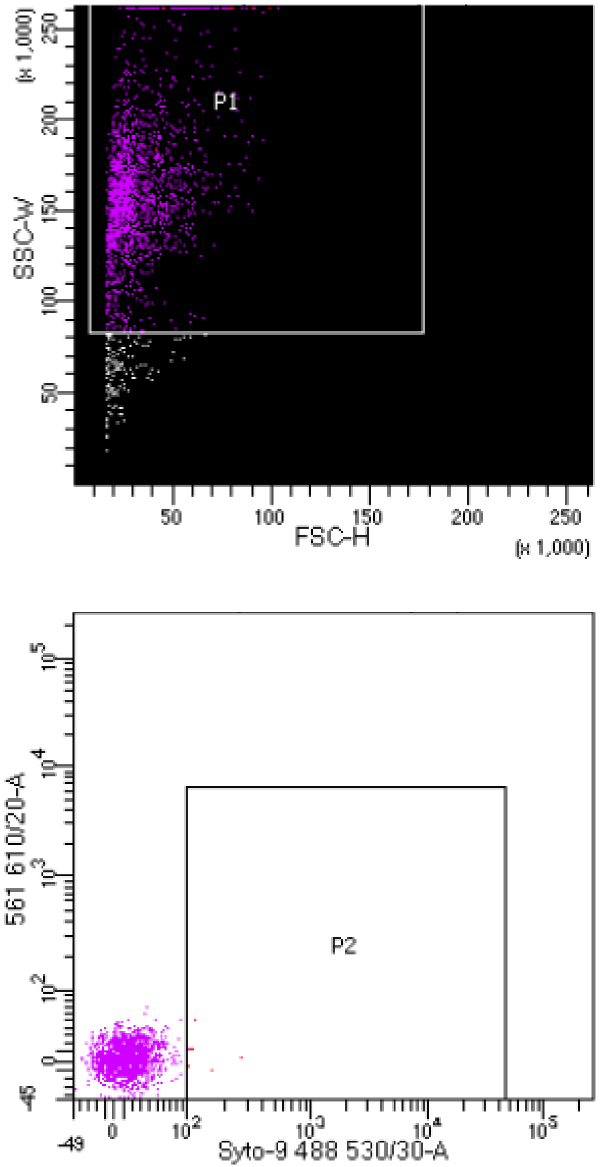


#### Capsules (medium)

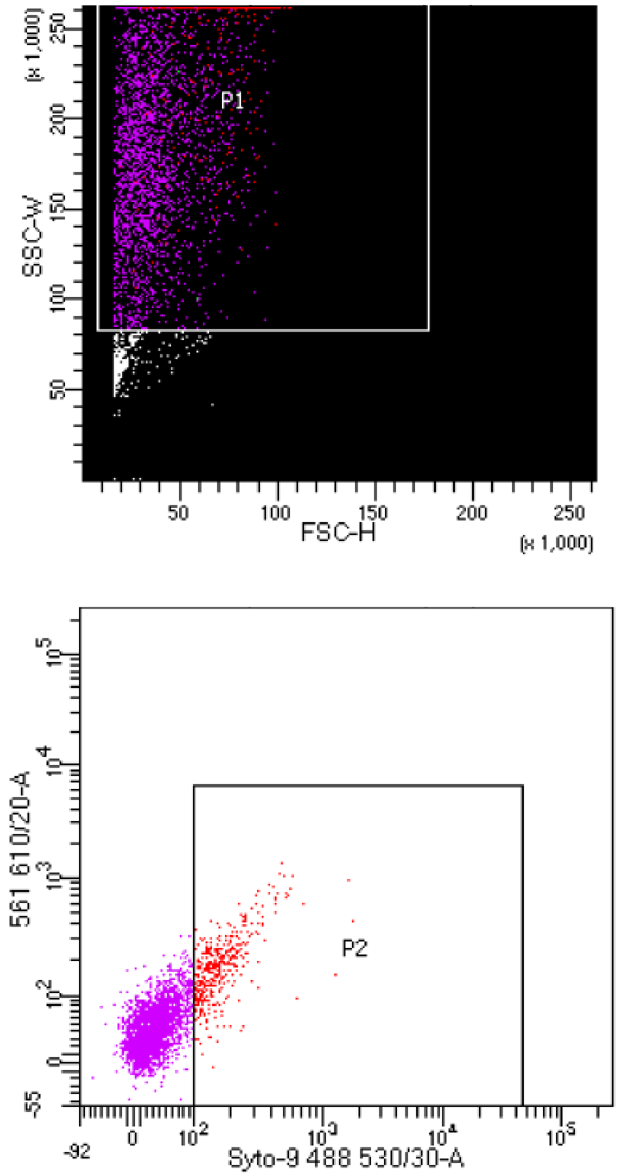

### Supplementary Figure 5

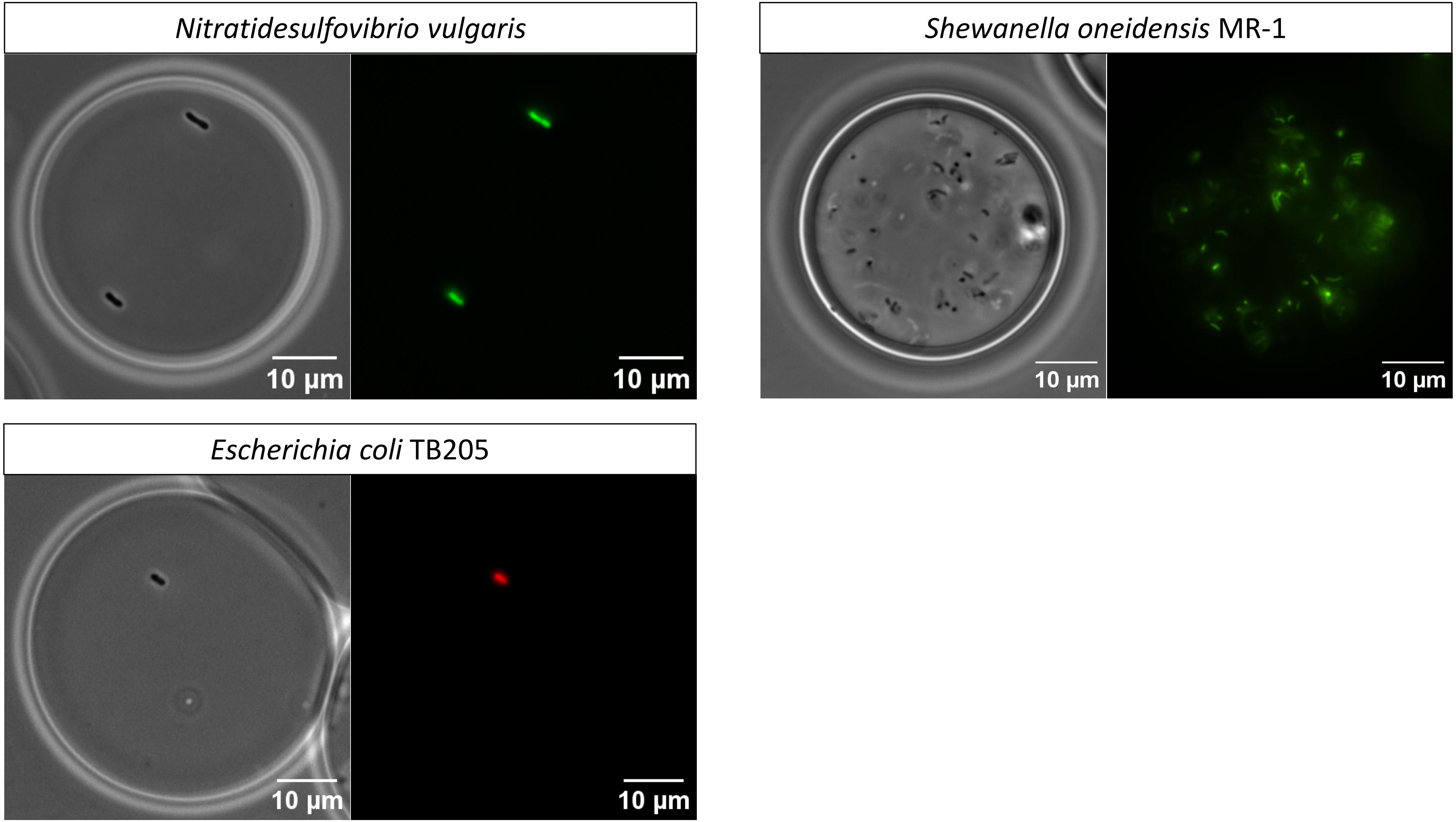

### Supplementary Figure 6

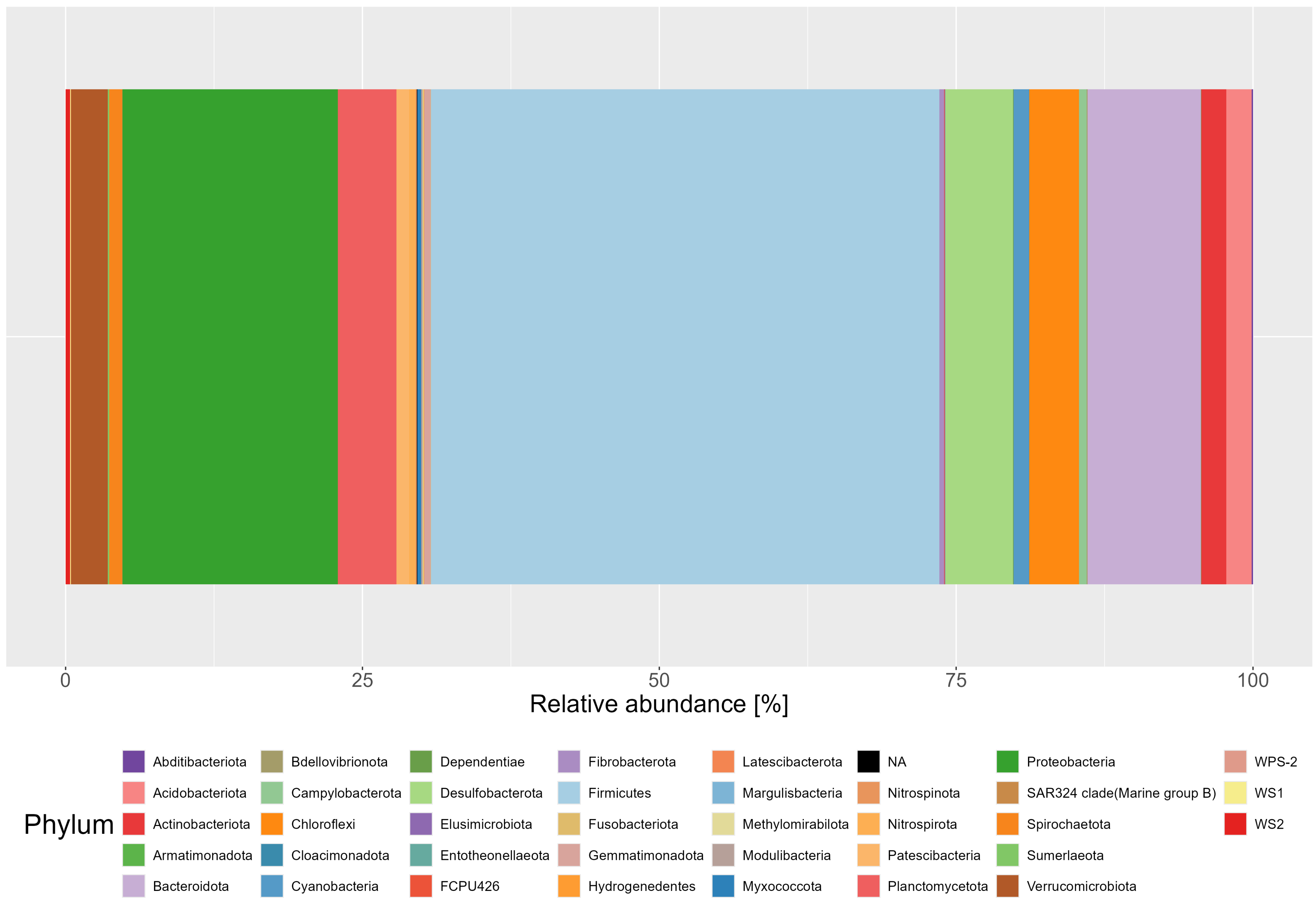

### Supplementary Figure 7

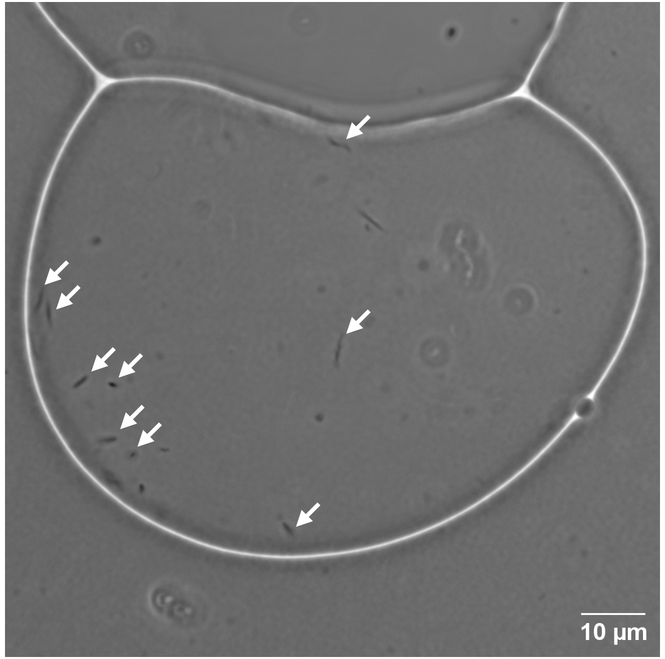


w-o droplets
15 h

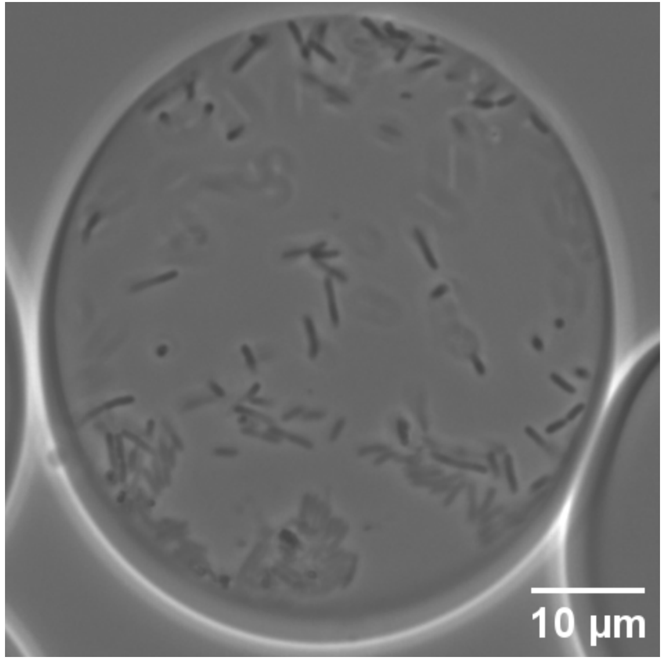


Capsules
15 h

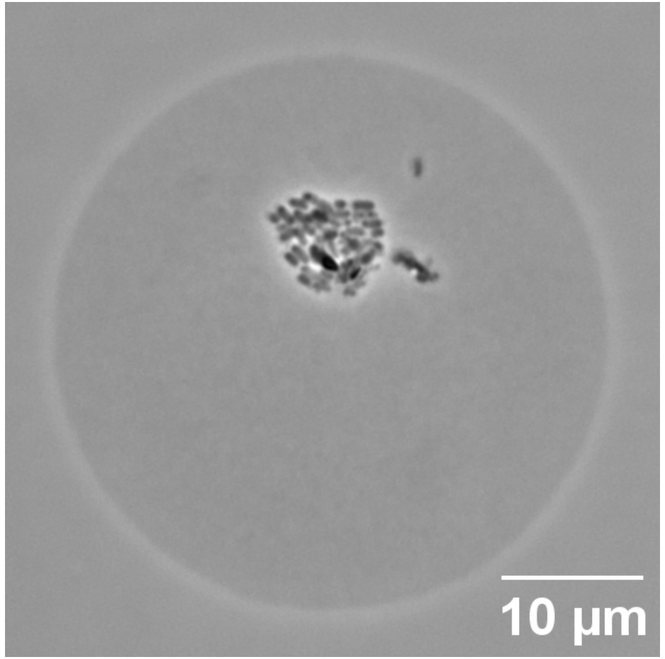


Agarose beads
4 d

### Supplementary Figure 8

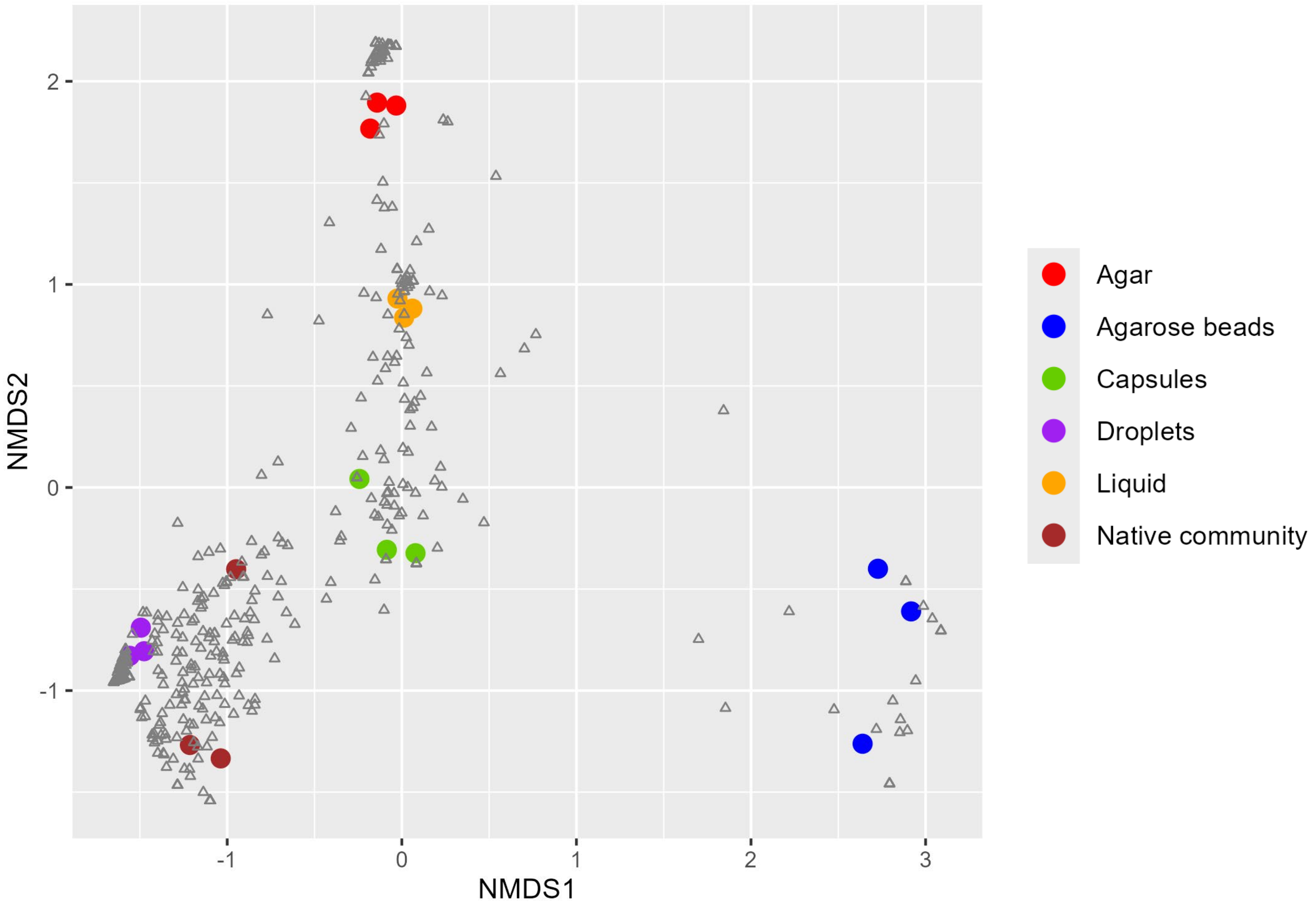

### Supplementary Figure 9

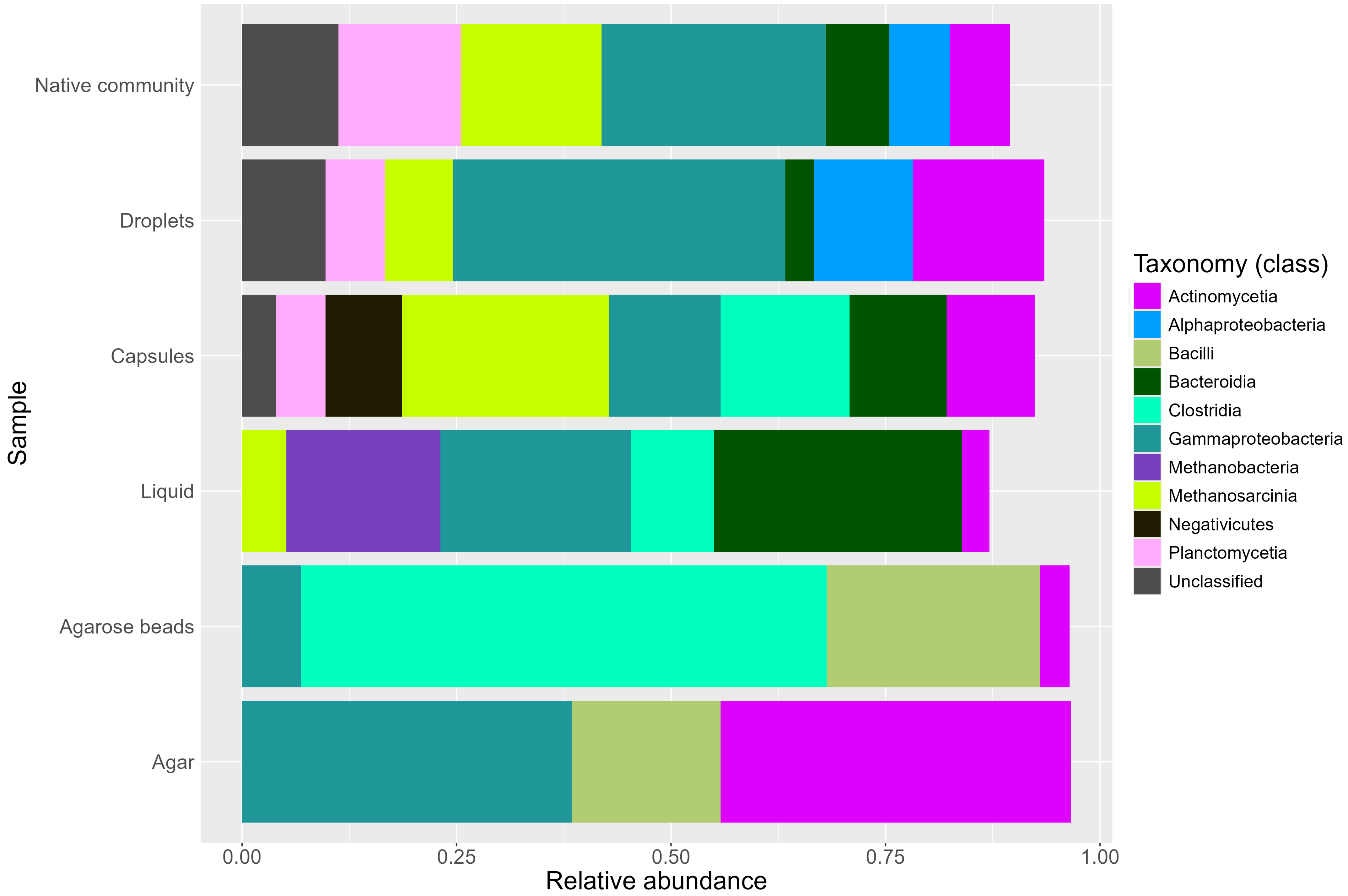

### Supplementary Figure 10

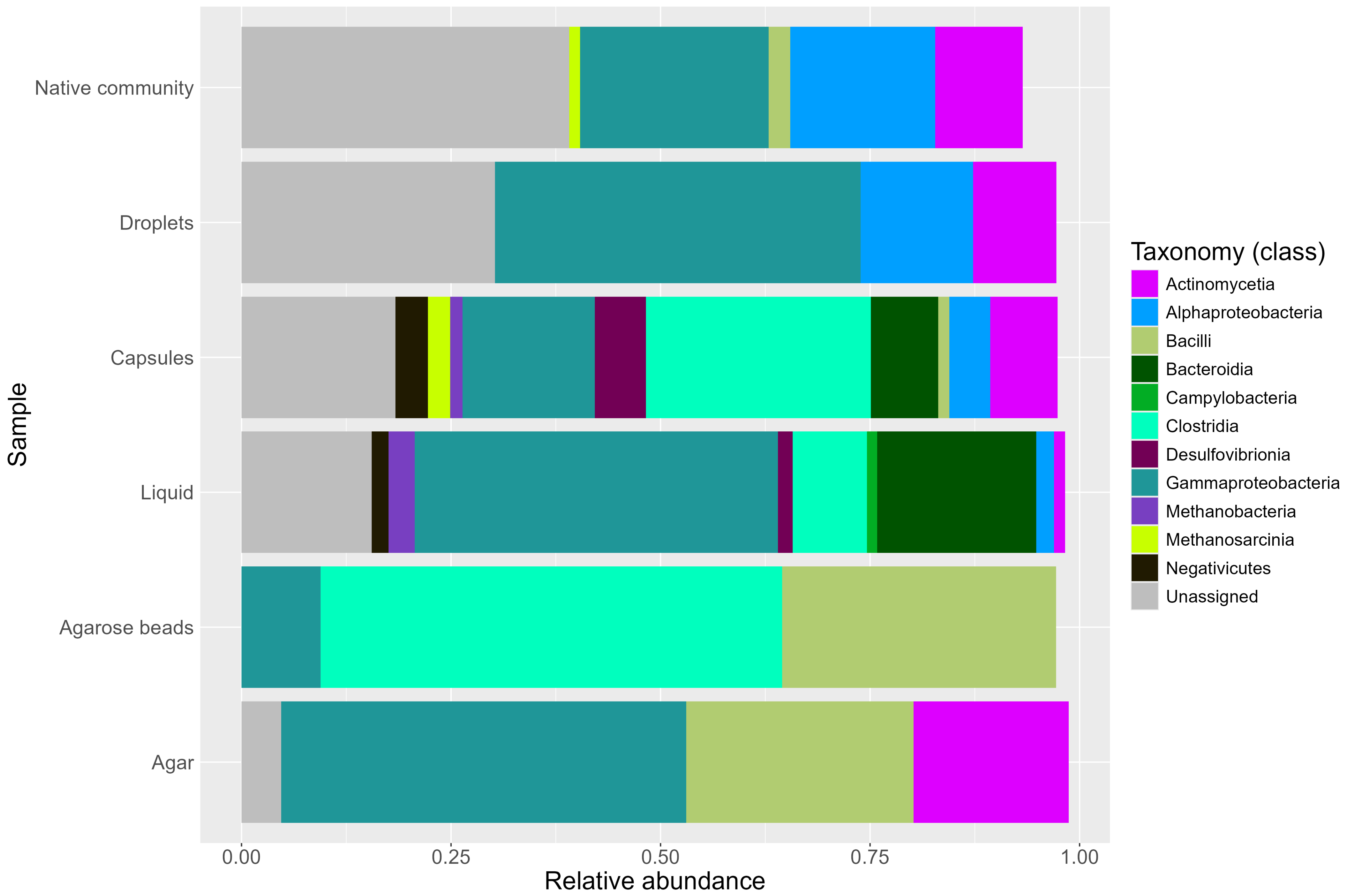

### Supplementary Figure 11

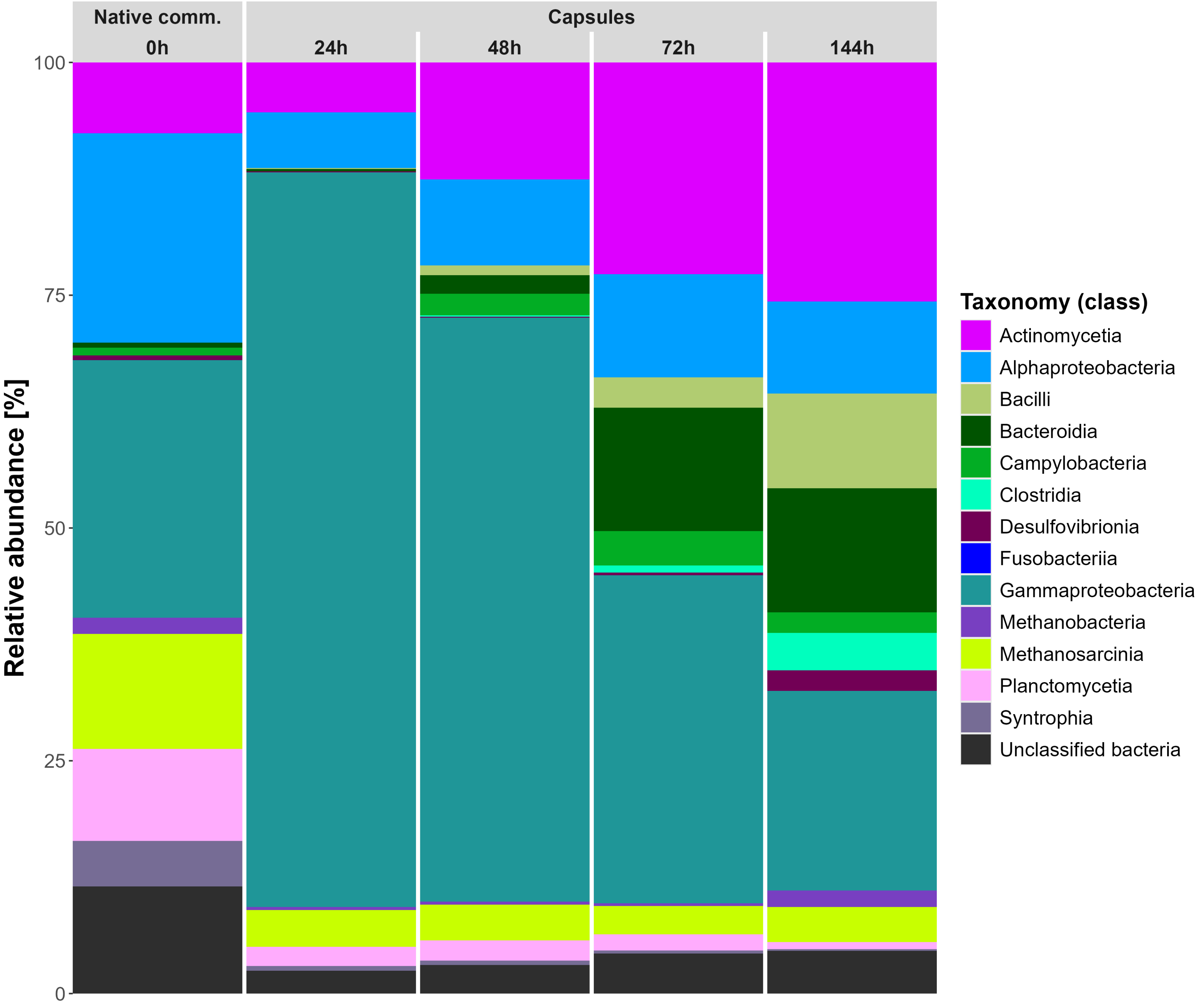

### Supplementary Figure 12

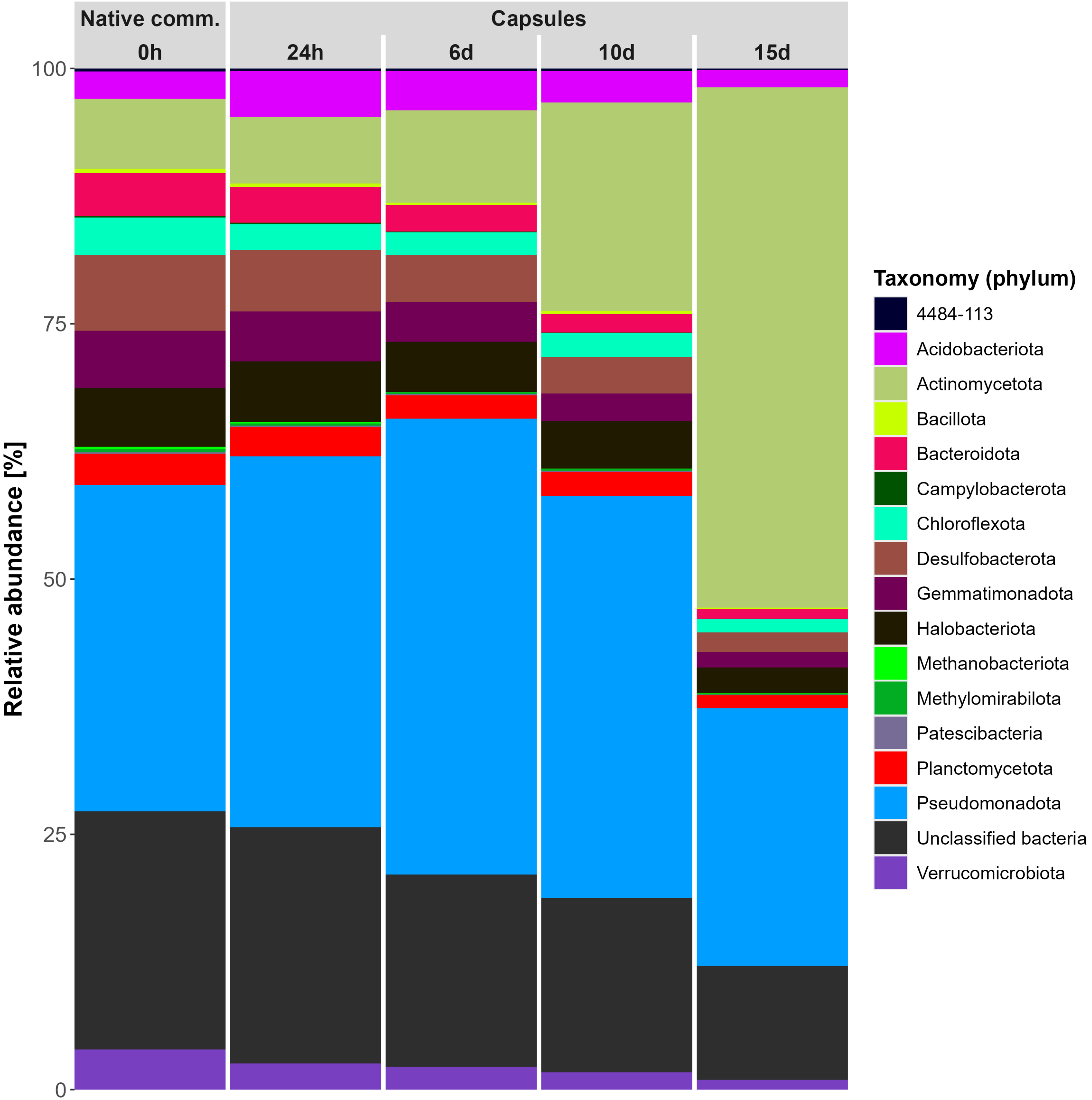

### Supplementary Figure 13

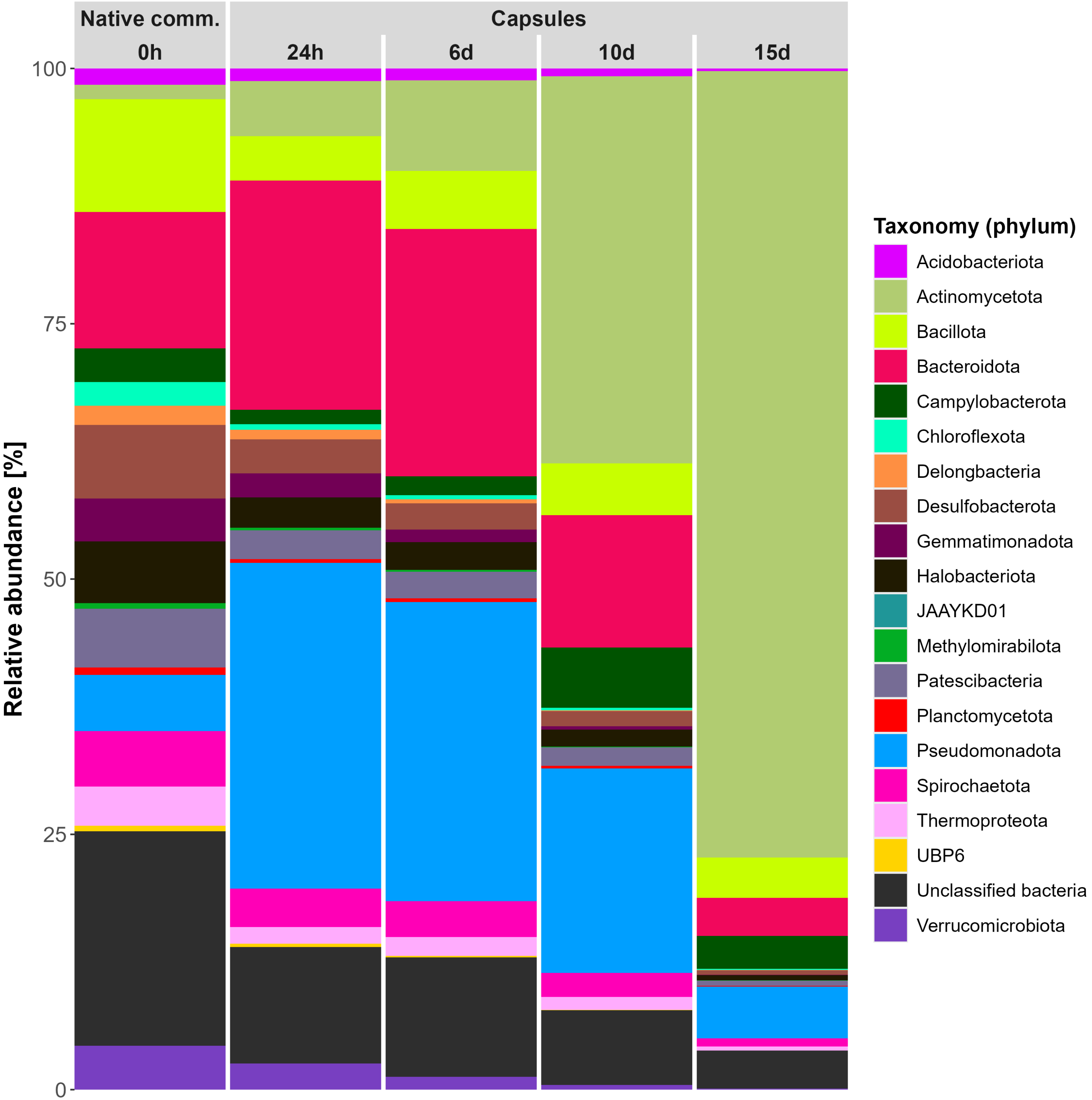

### Supplementary Figure 14

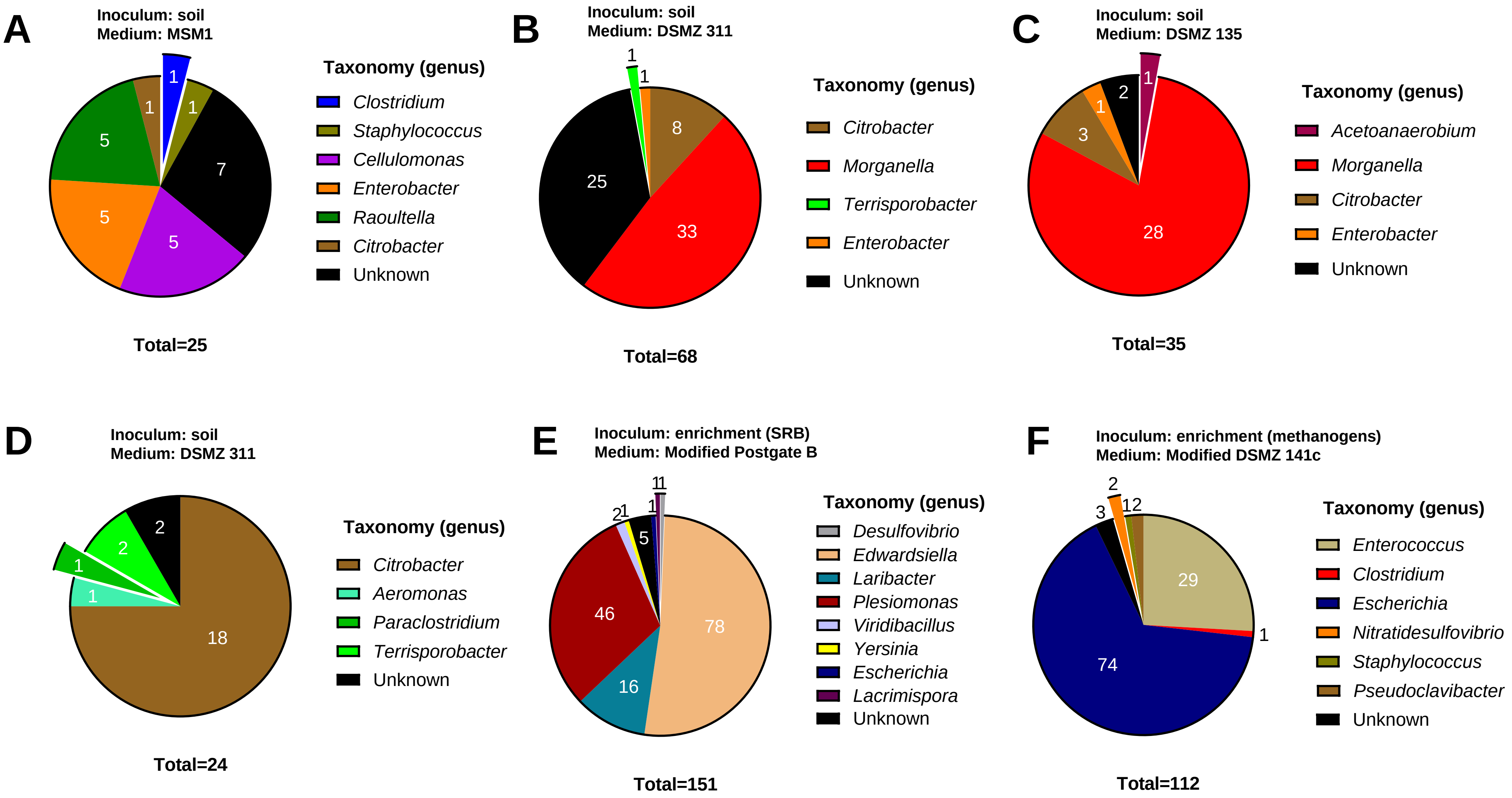

### Supplementary Figure 15

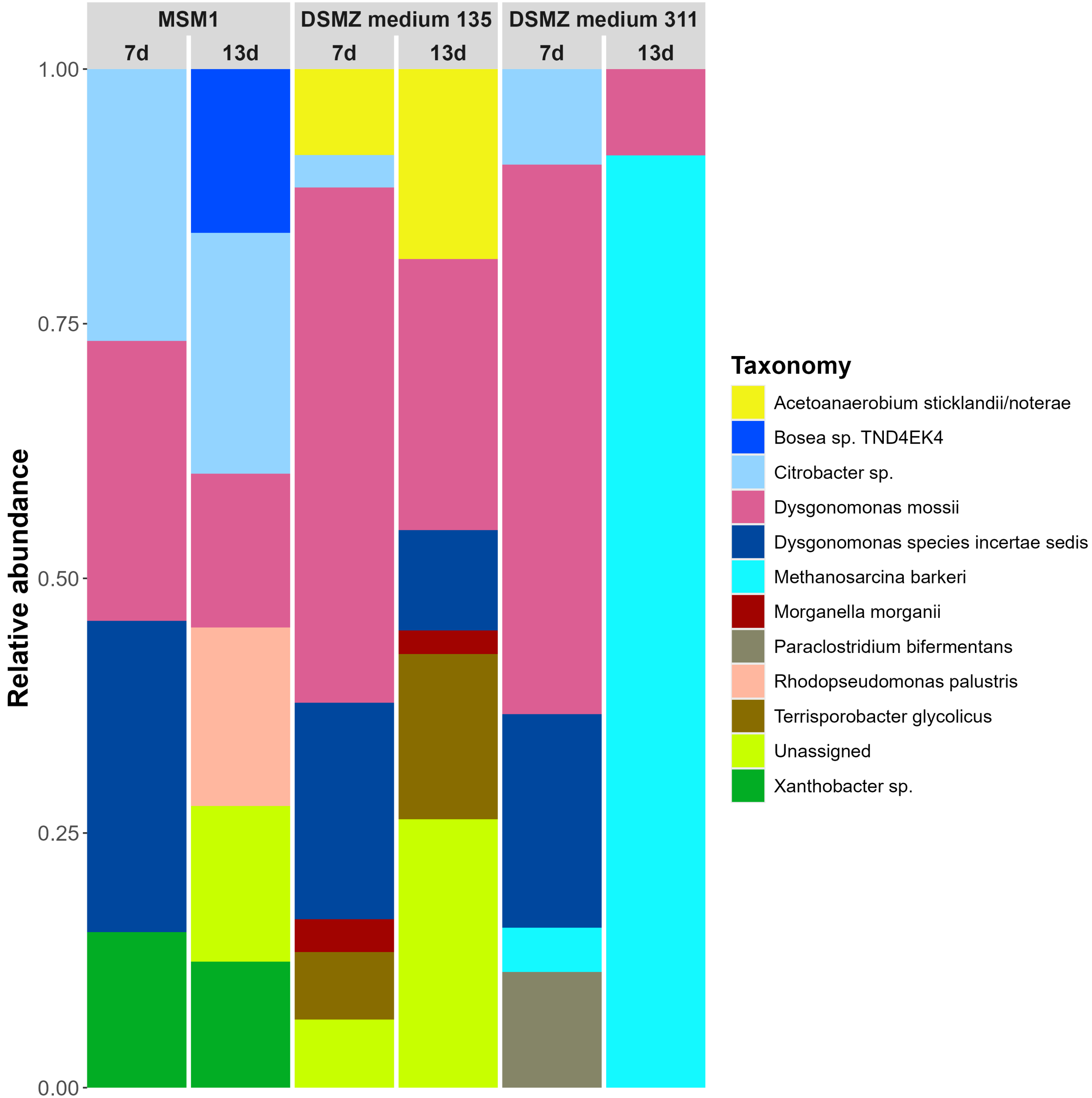

### Supplementary Figure 16

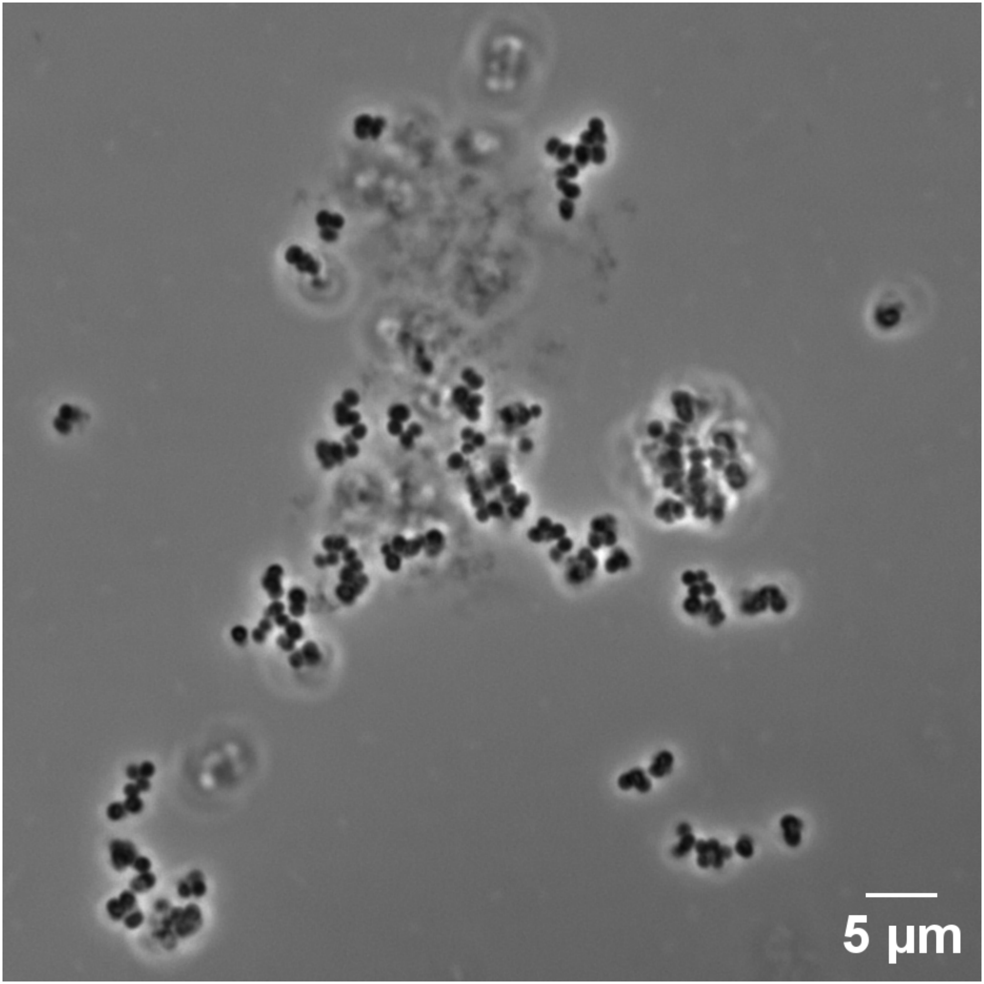

### Supplementary Figure 17

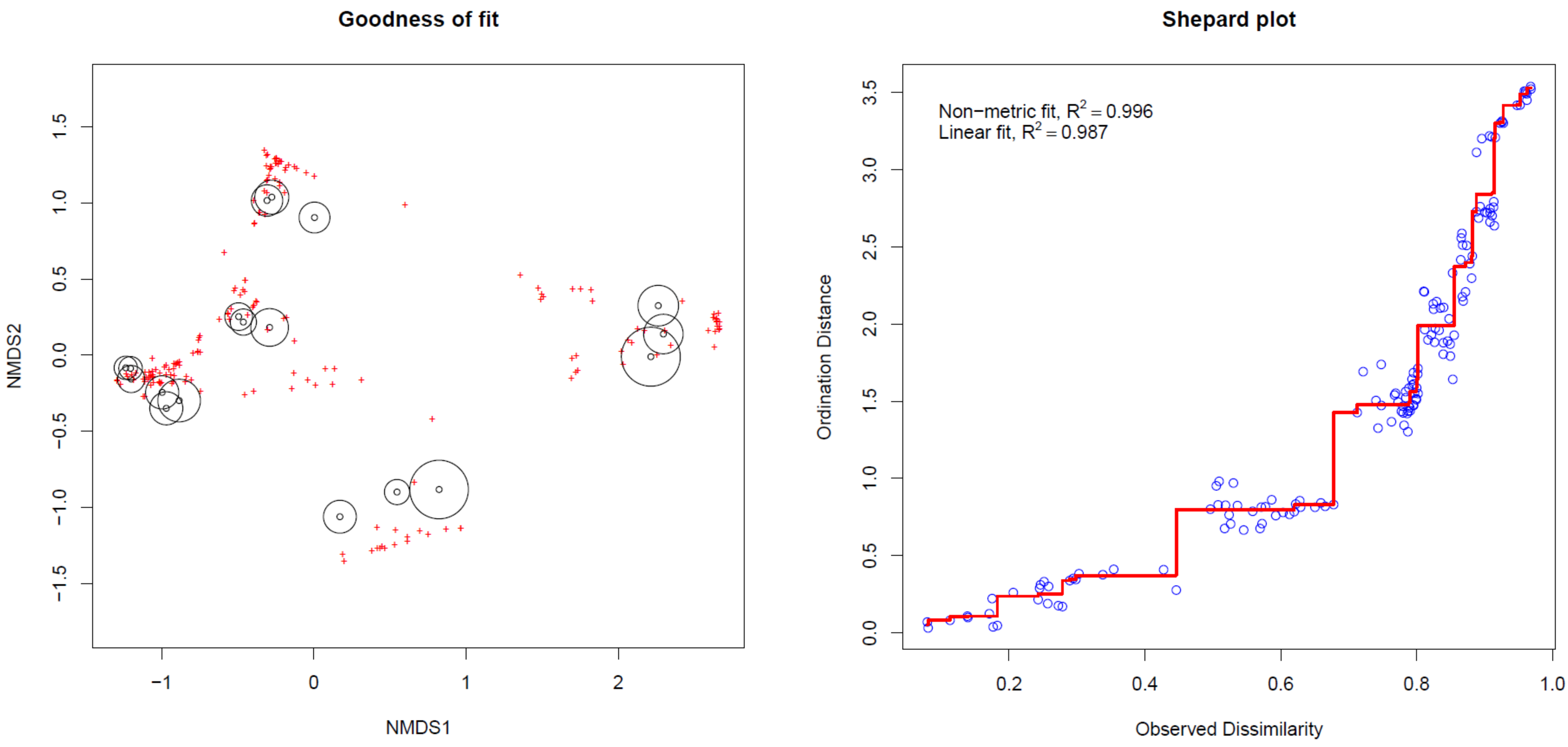

### Supplementary Figure 18

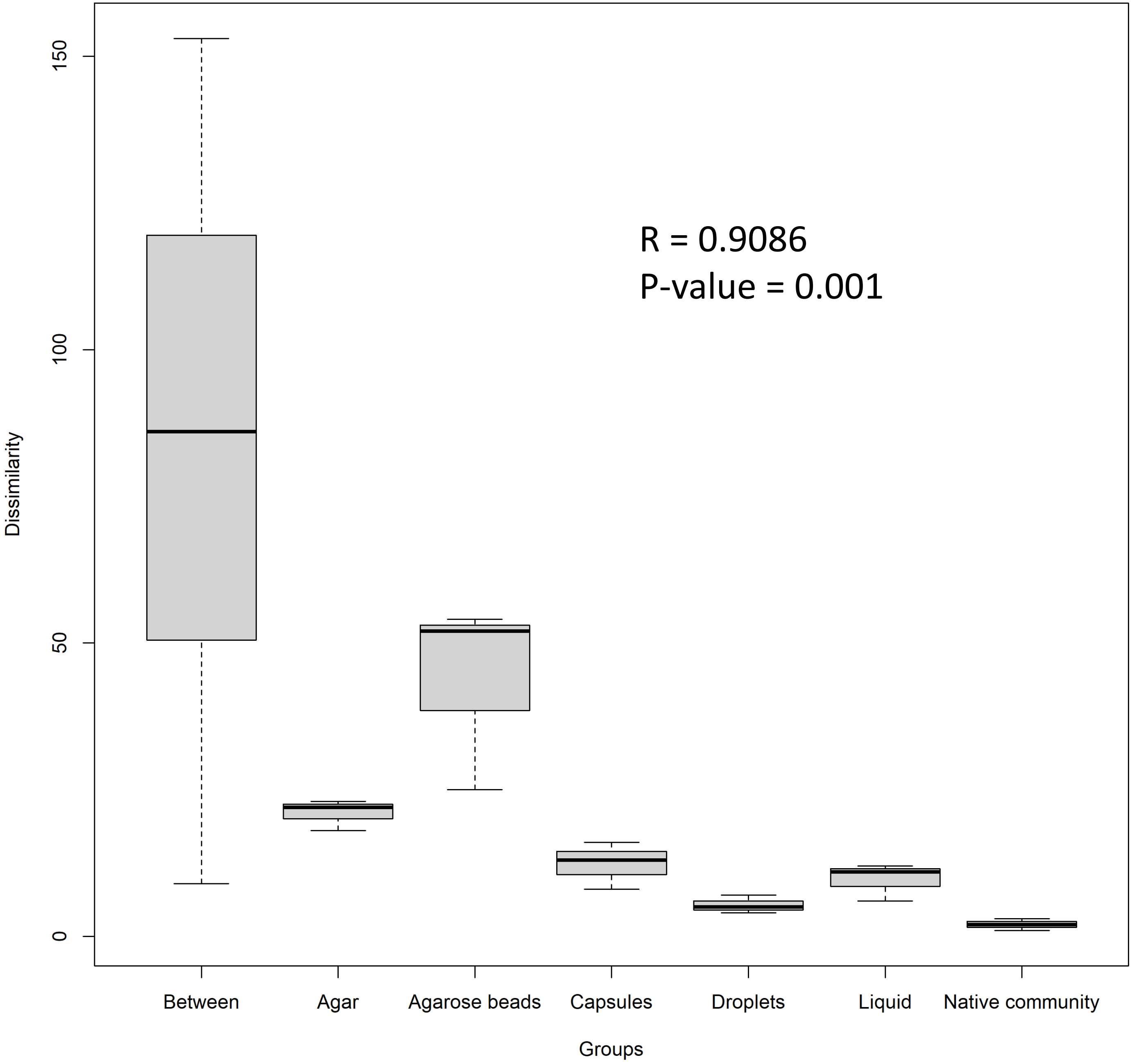
