## Supplementary Figure 4 for "High-throughput cultivation and isolation of environmental anaerobes using selectively permeable hydrogel capsules"

### Hydrogel capsule

- Permeable
- Liquid compartment
- Planktonic cells
- Compatible with FACS

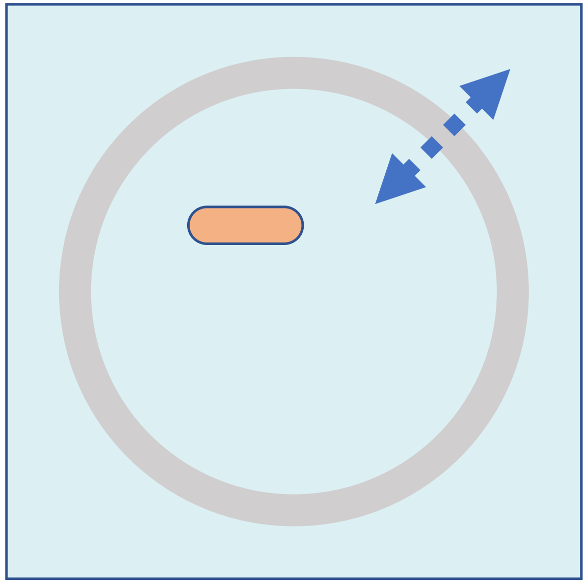

### Agarose bead

- Permeable
- Solid compartment (gel matrix)
- Sessile cells
- Compatible with FACS

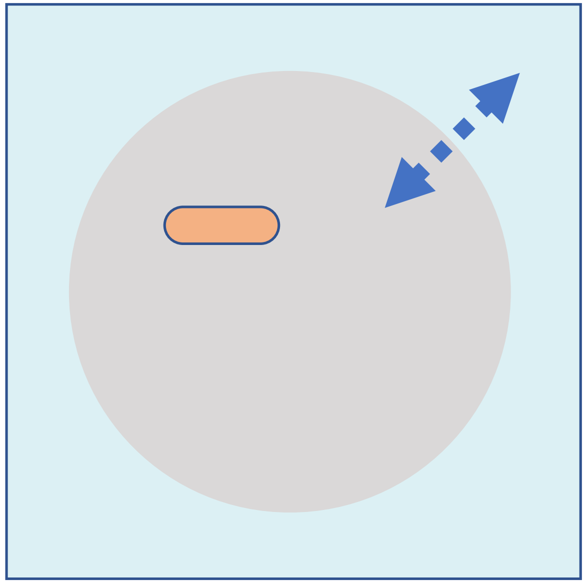

### Water-in-oil droplet

- Impermeable (water-oil barrier)
- Liquid compartment
- Planktonic cells
- Not directly compatible with FACS (need double emulsion)

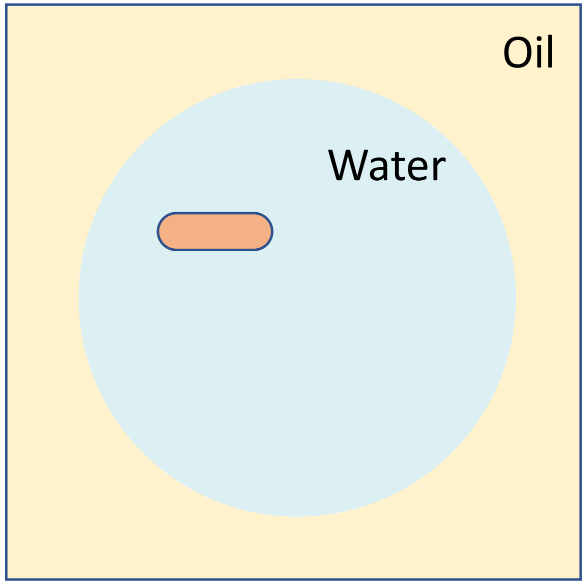
