## Supplementary Information for "High-throughput cultivation and isolation of environmental anaerobes using selectively permeable hydrogel capsules"

Media preparation

All media were brought to a boil, distributed in 200-ml serum bottles (100 ml / bottle) sealed with butyl rubber stoppers and aluminium crimps. The headspace was flushed with anoxic gases before autoclaving at 121°C for 30 min. The composition of the media used in the study can be found below:

Minimal soil medium (MSM): NaCl (1.0 g/l), MgCl_2_·6H_2_O (0.4 g/l), CaCl_2_·2H_2_O (0.1 g/l), NH_4_Cl (0.25 g/l), KH_2_PO_4_ (0.5 g/l), KCl (0.5 g/l), soil extract (10 ml/l for MSM1, 100 ml/l for MSM10), vitamins (cyanocobalamin (50 µg/l), thiamine (100 µg/l), 4-aminobenzoic acid (40 µg/l), D(+)-biotin (10 µg/l), nicotinic acid (100 µg/l), calcium D-(+)-pantothenate (50 µg/l), pyridoxine hydrochloride (150 µg/l)), pH 7.0, CO_2_/N_2_ (20/80 v/v) headspace. The soil extract was prepared by mixing dry soil (2-mm sieved) with Milli-Q water (1:2 w/w), autoclaving (121°C for 30 min) and filtering the supernatant (0.1 µm). An independent batch of MSM medium was prepared for each soil.

DSMZ 135 medium: NH₄Cl (1 g/l), KH₂PO₄ (0.33 g/l), K₂HPO₄ (0.45 g/l), MgSO₄·7H₂O (0.10 g/l), Modified Wolin's mineral solution (20 ml/l), Yeast extract (2 g/l), Sodium resazurin 0.1% w/v (0.5 ml/l), NaHCO₃ (10 g/l), D-Fructose (10 g/l), Wolin's 10x vitamin solution (1 ml/l), L-Cysteine HCl·H₂O (0.5 g/l), Na₂S·9H₂O (0.5 g/l).

DSMZ 311 medium: NH₄Cl (0.5 g/l), MgSO₄·7H₂O (0.5 g/l), CaCl₂·2H₂O (0.25 g/l), NaCl (2.25 g/l), FeSO₄·7H₂O 0.1% w/v in 0.1 N H₂SO₄ (2 ml/l), Trace element solution SL-10 (1 ml/l), Selenite-tungstate solution (1 ml/l), Yeast extract (2 g/l), Casitone (2 g/l), Betaine·H₂O (6.7 g/l), Sodium resazurin 0.1% w/v (0.5 ml/l), K₂HPO₄ (0.35 g/l), KH₂PO₄ (0.23 g/l), Na₂CO₃ (1 g/l), Wolin's vitamin solution (10x) (1 ml/l), L-Cysteine HCl·H₂O (0.3 g/l), Na₂S·9H₂O (0.3 g/l).

DSMZ 141c medium: KCl (0.34 g/l), MgCl₂·6H₂O (4 g/l), MgSO₄·7H₂O (3.45 g/l), NH₄Cl (0.25 g/l), CaCl₂·2H₂O (0.14 g/l), K₂HPO₄ (0.14 g/l), NaCl (18 g/l), Modified Wolin's mineral solution (10 ml/l), Fe(NH₄)₂(SO₄)₂·6H₂O 0.1% w/v (2 ml/l), Na-acetate (1 g/l), Yeast extract (2 g/l), Trypticase peptone (2 g/l), Sodium resazurin 0.1% w/v (0.5 ml/l), NaHCO₃ (5 g/l), Wolin's vitamin 10x solution (1 ml/l), L-Cysteine HCl·H₂O (0.5 g/l), Na₂S·9H₂O (0.5 g/l), Methanol (5 ml/l).

Modified DSMZ 141c medium: same recipe as DSMZ 141c medium, but with the following modifications: amendment of formate (0.3 g/l), no amendment of yeast extract, no amendment of trypticase peptone.

DSMZ medium 63: K₂HPO₄ 0.50 g, NH₄Cl 1.00 g, Na₂SO₄ 1.00 g, CaCl₂·2H₂O 0.10 g, MgSO₄·7H₂O 2.00 g, Na-DL-lactate 2.00 g, Yeast extract (1 g/l), Sodium resazurin 0.1% w/v (0.5 ml/l), FeSO₄·7H₂O (0.05 g/l), Na-thioglycolate (0.01 g/l), Ascorbic acid (0.01 g/l).

Modified Postgate B medium: as described by Kováč and Kushkevych [1], but with the following modifications: amendment of soil extract (10 ml/l) and sodium resazurin 0.1% w/v (0.5 ml/l).

Modified Wolin's mineral solution: Nitrilotriacetic acid (1.50 g/l), MgSO₄·7H₂O (3 g/l), MnSO₄·H₂O (0.5 g/l), NaCl (1 g/l), FeSO₄·7H₂O (0.1 g/l), CoSO₄·7H₂O (0.18 g/l), CaCl₂·2H₂O (0.1 g/l), ZnSO₄·7H₂O (0.18 g/l), CuSO₄·5H₂O (0.01 g/l), AlK(SO₄)₂·12H₂O (0.02 g/l), H₃BO₃ (0.01 g/l), Na₂MoO₄·2H₂O (0.01 g/l), NiCl₂·6H₂O (0.03 g/l), Na₂SeO₃·5H₂O (3.0 × 10⁻⁴ g/l), Na₂WO₄·2H₂O (4.0 × 10⁻⁴ g/l).

Selenite-tungstate solution: NaOH (0.5 g/l), Na₂SeO₃·5H₂O (3 mg/l), Na₂WO₄·2H₂O (4 mg/l).

Trace element solution SL-10: HCl (25%) (10 ml/l), FeCl₂·4H₂O (1.5 g/l), ZnCl₂ (70 mg/l), MnCl₂·4H₂O (100 mg/l), H₃BO₃ (6 mg/l), CoCl₂·6H₂O (190 mg/l), CuCl₂·2H₂O (2 mg/l), NiCl₂·6H₂O (24 mg/l), Na₂MoO₄·2H₂O (36 mg/l).

Wolin’s 10x vitamin solution: Biotin (20 mg/l), Folic acid (20 mg/l), Pyridoxine hydrochloride (100 mg/l), Thiamine HCl (50 mg/l), Riboflavin (50 mg/l), Nicotinic acid (50 mg/l), Calcium D-(+)-Pantothenate (50 mg/l), Vitamin B₁₂ (1 mg/l), p-Aminobenzoic acid (50 mg/l), (DL)-alpha-Lipoic acid (50 mg/l).

Enrichments of SRB and methanogens prior to encapsulation

SRB enrichment: 1 g of B_BH_ soil was inoculated in 10 ml modified Postgate B medium in a 25-ml Balch-type tube and incubated at 30°C. Consumption of sulfate and lactate was monitored with ion chromatography (Integrion HPIC, Thermo Fisher Scientific). After 10 days of incubation, the culture was encapsulated.

Methanogen enrichment: 1 g of B_BH_ soil was inoculated in 10 ml modified DSMZ 141c medium in a 25-ml Balch-type tube and incubated at 30°C. Consumption of H_2_ and CO_2_, as well as methane production, were confirmed by gas chromatography (456-GC, Scion Instruments). After 10 days, the culture was transferred to fresh medium (10% v/v inoculum) and the new culture was encapsulated after 8 days of incubation.

Bioinformatic analysis

MAGs were phylogenetically placed using GToTree (v1.8.4) with the 25 bacterial and archaeal marker genes HMM models. Functional annotation was performed using METABOLIC-G (v4.0) to identify metabolic pathways, while CoverM (v0.7.0) provided MAG abundance across samples after mapping with Strobealign (v0.12.0). The presence of a KEGG module was defined by the presence of 75% of its module steps. MAGs were assigned to aerobic taxa based on the presence of gene sets involved in the biosynthesis of cytochrome bc1 complex, cytochrome bd ubiquinol oxidase or cytochrome c oxidase, i.e., with any of the following KEGG modules present: M00151, M00152, M00153, M00154, M00155 or M00156.

MAGs were labelled depending on genome annotation. Those harbouring the genes involved in module M00496 (for reduction of sulfate to APS, reduction of APS to sulfite, and reduction of sulfite to sulfide) in the KEGG database were labelled as SRB. Those harbouring genes for any of the metabolic modules for methanogenesis (hydrogenotrophic, M00567; methylotrophic, M00564 or M00356; acetoclastic, M00357) were labelled as methanogens. MAGs comprising at least 5 of the 7 enzyme-encoding genes involved in the Wood-Ljungdahl pathway (module M00377) were labelled as acetogens.

In some instances, high-quality reads were used for mOTUs profiling with the default database (nr3.0.3) [2], in addition to the taxonomic classification with the GTDB-Tk database.

NMDS plots were produced with the metaMDS function (vegan package, R). Goodness of fit and Shepard plots were generated to check the validity and quality of the ordination (Fig. S17), while ANOSIM was used to confirm significance of the differences between groups (Fig. S18). Statistical analyses were performed using the Vegan package in R.

All analyses were conducted on the EPFL high-performance computing (HPC) cluster, utilizing SLURM (v23.11.10) and Apptainer (v1.2.5). Each node consisted of two Intel(R) Xeon(R) Platinum 8360Y processors running at 2.4 GHz, with 36 cores per processor (72 cores per node) and 3 TB of SSD storage.

FACS operating parameters

When sorting capsule, the sorter was equipped with a 100-µm nozzle. Sorting was performed at a frequency of 30 kHz, a pressure of 20 psi and an event rate of 200 s^-1^.

Table S1: **soil physical and chemical parameters**

| **Soil** | **A_AH_** | **B_AH_** | **A_BH_** | **B_BH_** |
| --- | --- | --- | --- | --- |
| Clay content (%) | 25.01 | 26.64 | 20.96 | 26.67 |
| Silt content (%) | 38.15 | 57.93 | 42.20 | 39.83 |
| Sand content (%) | 36.84 | 15.42 | 36.84 | 33.50 |
| Texture | loam | silt loam | loam | loam |
| pH | 7.92 | 7.49 | - | - |
| TOC (wt %) | 2.99 | 7.71 | - | - |
| C (wt.%)^*^ | 4.78 ± 0.04 | 8.04 ± 0.07 | - | - |
| H (wt.%) ^*^ | 0.692 ± 0.00 | 1.32 ± 0.01 | - | - |
| N (wt.%)^*^ | 0.25 ± 0.00 | 0.6 ± 0.01 | - | - |
| S (wt.%)^*^ | 0.070 ± 0.02 | 0.10 ± 0.00 | - | - |
| Ca [mg/kg] | 50036 | 19427 | - | - |
| Fe [mg/kg] | 17960 | 28707 | - | - |
| K [mg/kg] | 9116 | 11863 | - | - |
| Mg [mg/kg] | 7473 | 9779 | - | - |
| Mn [mg/kg] | <loq^**^ | 130 | - | - |
| Na [mg/kg] | 2168 | 1906 | - | - |
| P [mg/kg] | <loq^**^ | 801 | - | - |
| As [mg/kg] | 0.629 | 0.893 | - | - |

^*^Results are indicated as means of three analyses ± standard deviation.

^**^limit of quantification (loq) = 0.5 mg/l (ICP-OES).

- Indicates no data collected.

**Movie legends**

Movie 1

**Microfluidic generation of hydrogel capsules**. The time lapse shows the junction in the microfluidic chip where the aqueous phase (mix of two polymer components) meets the oil phase, resulting in an emulsion.

Movie 2

**Time lapse of *Nitratidesulfovibrio vulgaris* growing within hydrogel capsules (2-day incubation)**.

Movie 3

**Time lapse of *E. coli* TB205** **growing within hydrogel capsules (24 h incubation)**.

Movie 4

**Time lapse of *Paraclostridium bifermentans* EML growing within hydrogel capsules (24 h incubation)**.

Movie 5

**Time lapse of *Thermoanaerobacter kivui* growing within hydrogel capsules (24 h incubation)**.

Movie 6

**Time lapse of *Shewanella oneidensis* MR-1 growing within hydrogel capsules (24 h incubation)**.

**References**

1. Kováč J, Kushkevych I. New modification of cultivation medium for isolation and growth of intestinal sulfate-reducing bacteria. 2017.

2. Ruscheweyh HJ, Milanese A, Paoli L *et al.* Cultivation-independent genomes greatly expand taxonomic-profiling capabilities of mOTUs across various environments. *Microbiome* 2022;**10**:1–12.
